# *In Vivo* Brain Glutathione is Higher in Older Age and Correlates with Mobility

**DOI:** 10.1101/2020.10.14.339507

**Authors:** K. E. Hupfeld, H. W. Hyatt, P. Alvarez Jerez, M. Mikkelsen, C. J. Hass, R. A. E. Edden, R. D. Seidler, E. C. Porges

**Affiliations:** Department of Applied Physiology and Kinesiology, University of Florida, Gainesville, FL; Russell H. Morgan Department of Radiology and Radiological Science, The Johns Hopkins University School of Medicine, Baltimore, MD; F. M. Kirby Research Center for Functional Brain Imaging, Kennedy Krieger Institute, Baltimore, MD; Department of Neurology, University of Florida, Gainesville, FL; Department of Clinical and Health Psychology, University of Florida, Gainesville, FL

**Keywords:** glutathione (GSH), magnetic resonance spectroscopy (MRS), Hadamard Encoding and Reconstruction of MEGA-Edited Spectroscopy (HERMES), aging, balance, gait, manual dexterity

## Abstract

Brain markers of oxidative damage increase with advancing age. In response, brain antioxidant levels may also increase with age, although this has not been well investigated. Here we used edited magnetic resonance spectroscopy to quantify endogenous levels of glutathione (GSH, one of the most abundant brain antioxidants) in 37 young (mean: 21.8 (2.5) years; 19 F) and 23 older adults (mean: 72.8 (8.9) years; 19 F). Accounting for age-related atrophy, we identified higher frontal and sensorimotor GSH levels for the older compared to the younger adults. For the older adults only, higher sensorimotor (but not frontal) GSH was correlated with poorer balance, gait, and manual dexterity. This suggests a regionally-specific relationship between higher brain oxidative stress levels and motor performance declines with age. We suggest these findings reflect a compensatory upregulation of GSH in response to increasing brain oxidative stress with normal aging. Together, these results provide insight into age differences in brain antioxidant levels and implications for motor function.

## Introduction

The role of oxidative stress in brain aging has been studied since the emergence of the free radical theory of aging. This theory posits that the cumulative result of a lifetime of oxidative insult is diminished tissue functioning and the aging phenotype (Harman, 1955). While evidence exists both for and against the free radical theory of aging, the literature largely agrees that markers of brain oxidative damage increase with advancing age (Chakrabarti et al., 2011). The brain is a highly oxidative organ that consumes 20% of the body’s total oxygen uptake despite accounting for only 2% of the body’s total weight (Hyder et al., 2013; Quastel & Wheatley, 1932). This high rate of oxygen consumption, along with high levels of oxidizable iron molecules and polyunsaturated fats, increases the propensity of the brain to form reactive oxygen species (ROS). ROS production is a natural phenomenon that contributes to cell signaling. Excessive ROS production can lead to oxidative damage and requires detoxification of ROS molecules by antioxidant sources to prevent oxidative stress. Therefore, it is important to understand if antioxidant levels change in the brain with aging and if these changes relate to functional impairments, such as declines in cognition and motor control.

Glutathione (GSH) is one of the most abundant antioxidant sources in the central nervous system and plays a key role in the maintenance of redox homeostasis (Rice et al., 2002). Within the brain, GSH abundance appears to vary by cell type (Huang & Philbert, 1995; Langeveld et al., 1996; Raps et al., 1989; Rice & Russo-Menna, 1997) and brain region (Calabrese et al., 2002; Nezhad et al., 2017; Perry et al., 1971; Srinivasan et al., 2010). For detailed reviews of GSH biochemical characteristics, functions, and locations, see (Dringen, 2000; Dwivedi et al., 2020; Rae & Williams, 2017). While several studies have measured age differences in cortical GSH, current understanding of such changes remains equivocal. Rodent model studies have suggested that GSH decreases with age (e.g., (Chen et al., 1989; Liu, 2002; Sasaki et al., 2001)), but others have found no changes (e.g., (Asuncion et al., 1996; Hussain et al., 1995)). Post-mortem human work has reported no age-related differences in brain GSH levels (Tong et al., 2016; Venkateshappa et al., 2012), lower GSH levels among older adults (Venkateshappa et al., 2012), and similar or higher GSH levels in older age (Tong et al., 2016). Of note, these previous attempts to measure GSH levels with aging have been hampered by a lack of non-invasive procedures; furthermore, measurements in post-mortem conditions are subject to GSH breakdown (Perry et al., 1981), complicating interpretations and comparison to *in vivo* GSH levels.

Recent advances in spectral editing now make it possible to resolve GSH with magnetic resonance spectroscopy (MRS) (Saleh et al., 2016). Previously, without the use of spectral editing, GSH could not be quantified at 3 Tesla (Nezhad et al., 2017). Only one study to date has used edited MRS to compare GSH levels between normal young and older adults (Emir et al., 2011). This study scanned the occipital cortex and reported that GSH levels were 30% lower for older compared to younger adults (Emir et al., 2011). In the present study, we used Hadamard Encoding and Reconstruction of MEGA-Edited Spectroscopy (HERMES) (Saleh et al., 2016; Saleh et al., 2019) to examine age differences in GSH levels in the frontal and sensorimotor cortices, brain regions involved in cognitive function and mobility, respectively. Of note, as a recent review (Cleeland et al., 2019) discusses, few studies have explored age-related changes in neurometabolite levels in the sensorimotor cortex, and no previous studies have characterized age differences in GSH levels within the sensorimotor cortex.

While our current understanding of how aging affects brain GSH levels is limited, some evidence suggests that cortical GSH may be associated with cognitive and sensorimotor function. Normal aging results in cognitive decrement (Anstey & Low, 2004), as well as widespread motor decline, including difficulties with fine motor control (Seidler & Stelmach, 1995), balance (Downs et al., 2014), and walking (Rantakokko et al., 2013). Past work has found lower brain GSH levels in patients with mild cognitive impairment (MCI) and Alzheimer’s disease (AD) compared to normal aging (Mandal et al., 2015; Mandal et al., 2012). Lower levels of brain GSH in the frontal cortex (Mandal et al., 2015; Mandal et al., 2012), parietal cortex (Oeltzschner et al., 2019), and hippocampus (Mandal et al., 2015) have been associated with a larger degree of cognitive impairment in MCI and AD. Despite this, the limited work in normal aging has not found relationships between MRS-measured brain GSH levels and cognitive status (Chiang et al., 2017; Emir et al., 2011). Moreover, relationships between motor function and brain GSH levels have not yet been tested for normal older adults, although there is some support for a relationship between GSH and motor function, given that GSH levels are altered in various movement disorders. For instance, MRS-measured GSH levels are decreased in multiple sclerosis (motor cortex (Srinivasan et al., 2010), frontal cortex (Choi et al., 2011)), amyotrophic lateral sclerosis (motor cortex (Weerasekera et al., 2019; Weiduschat et al., 2014)), and spinocerebellar ataxia (cerebellum (Doss et al., 2015)). Although these studies did not report relationships between brain GSH levels and motor performance or disease severity, past rodent work has found that transient basal ganglia GSH depletion results in pronounced sensorimotor impairments (Díaz-Hung et al., 2014). Taken together, it is plausible that alterations in regional brain GSH levels may affect cognitive and sensorimotor function, although it is unclear whether this relationship would be evident in normal aging, or only in pathological conditions. In the present work, we tested associations between brain GSH levels and performance. We predicted regionally-specific relationships in which frontal GSH levels would be associated with cognitive performance, and sensorimotor GSH levels would be associated with motor performance.

Overall, it remains unclear how human brain GSH levels alter with aging and whether brain GSH is associated with cognitive or motor function. Based on the limited *in vivo* human work (Emir et al., 2011) and the larger body of animal and post-mortem human studies, it is plausible that brain GSH levels would be lower in older adults. If GSH levels are lower in older age, this may indicate greater oxidative stress burden, thereby exhausting brain antioxidant capacity. However, if GSH levels are higher for older adults, this could suggest a compensatory upregulation response to increased oxidative stress. For instance, there is evidence that mild stress increases brain GSH levels; this upregulation of GSH is thought to provide protection against more severe oxidative stress (for review, see (Maher, 2005)). Thus, it is possible that normal aging could be associated with higher brain GSH levels as a compensatory response to generalized aging processes.

The aims of the present study included: 1) to determine whether there are age differences in *in vivo* MRS-measured brain GSH levels in the frontal and sensorimotor cortices; 2) to characterize regional differences in brain GSH levels; and 3) to characterize the relationships between brain GSH levels and cognitive and motor function.

## Results

37 young and 23 older adults completed cognitive and motor testing, as well as collection of MRS data from voxels placed in the frontal and sensorimotor cortices. Of note, we applied the Benjamini-Hochberg false discovery rate (FDR) correction (Benjamini & Hochberg, 1995) to all *p*-values reported below; aside from two cases where the corrected *p*-values were *p* < 0.10, all results remained significant (*p* < 0.05) after applying this correction for multiple comparisons.

### Demographics

There were no significant age differences for most demographic variables, including sex, alcohol use, handedness, or footedness. Importantly, there were also no age differences in the number of days elapsed between the two testing sessions or in the difference in start time for the two sessions. See Table A1 for complete demographic information.

### Higher GSH Levels in Older Age

The older adult group exhibited cortical atrophy, with both voxels showing a lower gray matter fraction and higher cerebrospinal fluid (CSF) fraction compared to the younger adults (Table A2; Fig. 1). Older adults also had less white matter within the frontal voxel compared to younger adults. Older adults had significantly higher CSF-corrected GSH levels in both voxels (Fig. 2; although age differences in frontal GSH levels remained only at trend-level significance, *p* = 0.066, after FDR correction for multiple comparisons). This difference in CSF-corrected GSH levels implies that there is an age-related increase in cortical GSH concentration within the tissue that remains in the voxel after accounting for age-related atrophy. Importantly, there was no age difference in GSH fit error or water full width at half maximum (FWHM).

**Fig. 1.**
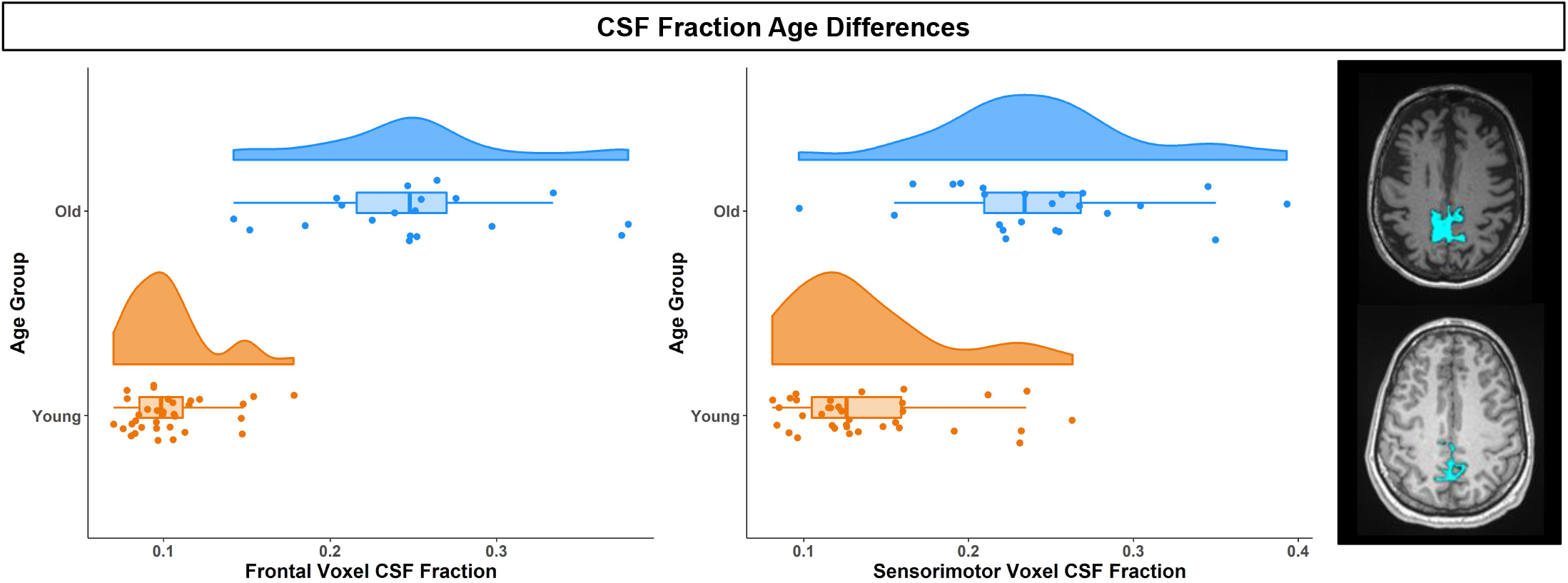
Higher CSF Fraction in Older Age. *Left*. CSF fraction within the frontal (left) and sensorimotor (right) voxels for older (blue) and young (orange) adults. In both voxels, older adults had higher CSF concentrations compared to young adults. *Right*. CSF fraction (blue) within the sensorimotor voxel, shown for a single older (top) and a single younger (bottom) participant. The CSF fraction is overlaid onto each subject’s native space *T*_1_-weighted anatomical image.

**Fig. 2.**
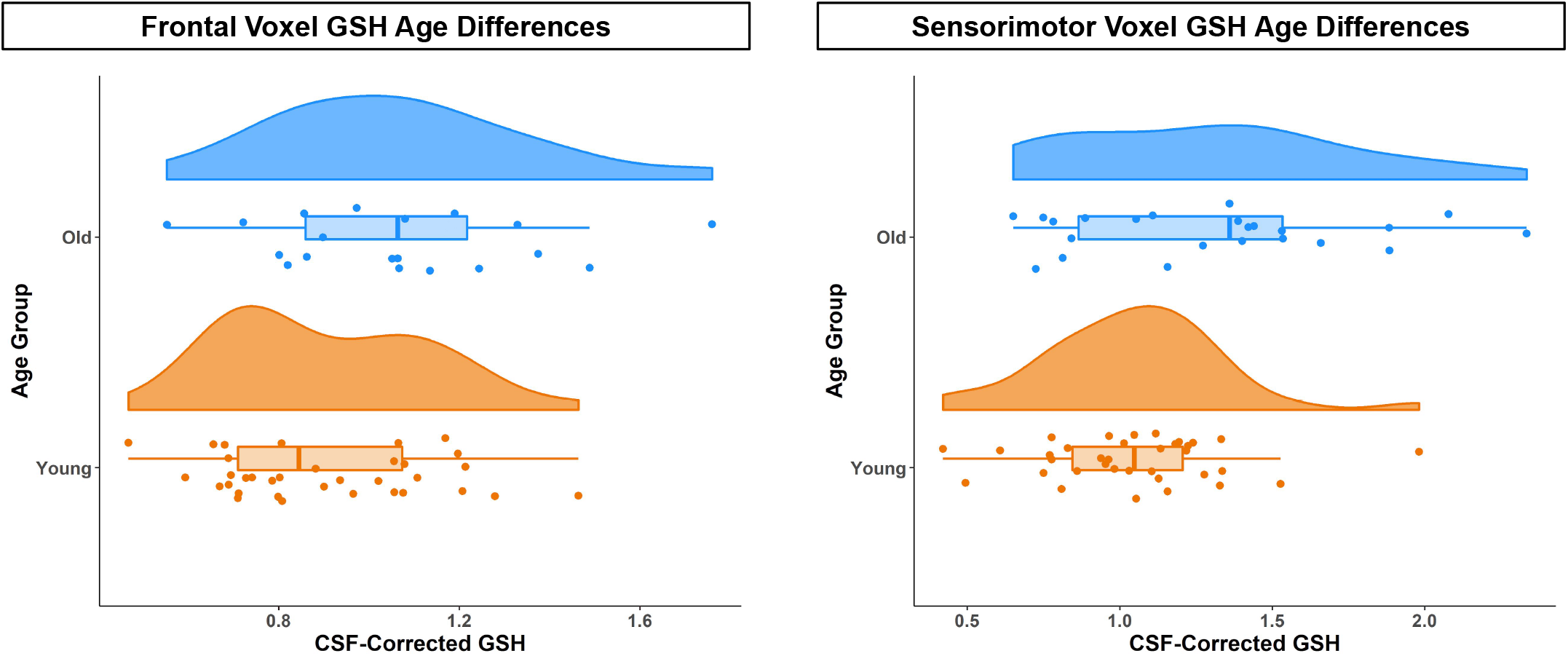
Higher GSH Levels in Older Age. CSF-corrected GSH levels for older (blue) and young (orange) adults in the frontal (left) and sensorimotor (right) voxels. Across both voxels, older adults had higher CSF-corrected GSH levels.

### Higher GSH Levels in Sensorimotor versus Frontal Cortex

Frontal GSH levels positively correlated with sensorimotor GSH levels for both the young and older adults (Table A3; Fig. 3). This relationship was in the same direction for both groups, and the correlation strength did not significantly differ by age. Within subjects, both groups also had higher GSH levels in the sensorimotor voxel compared to the frontal voxel. The magnitude of this regional effect did not differ between the age groups. For young adults, the gray matter and CSF fraction was higher and the white matter fraction was lower in the sensorimotor compared to the frontal voxel. However, for older adults, there were no significant differences in tissue composition between the two voxels.

**Fig. 3.**
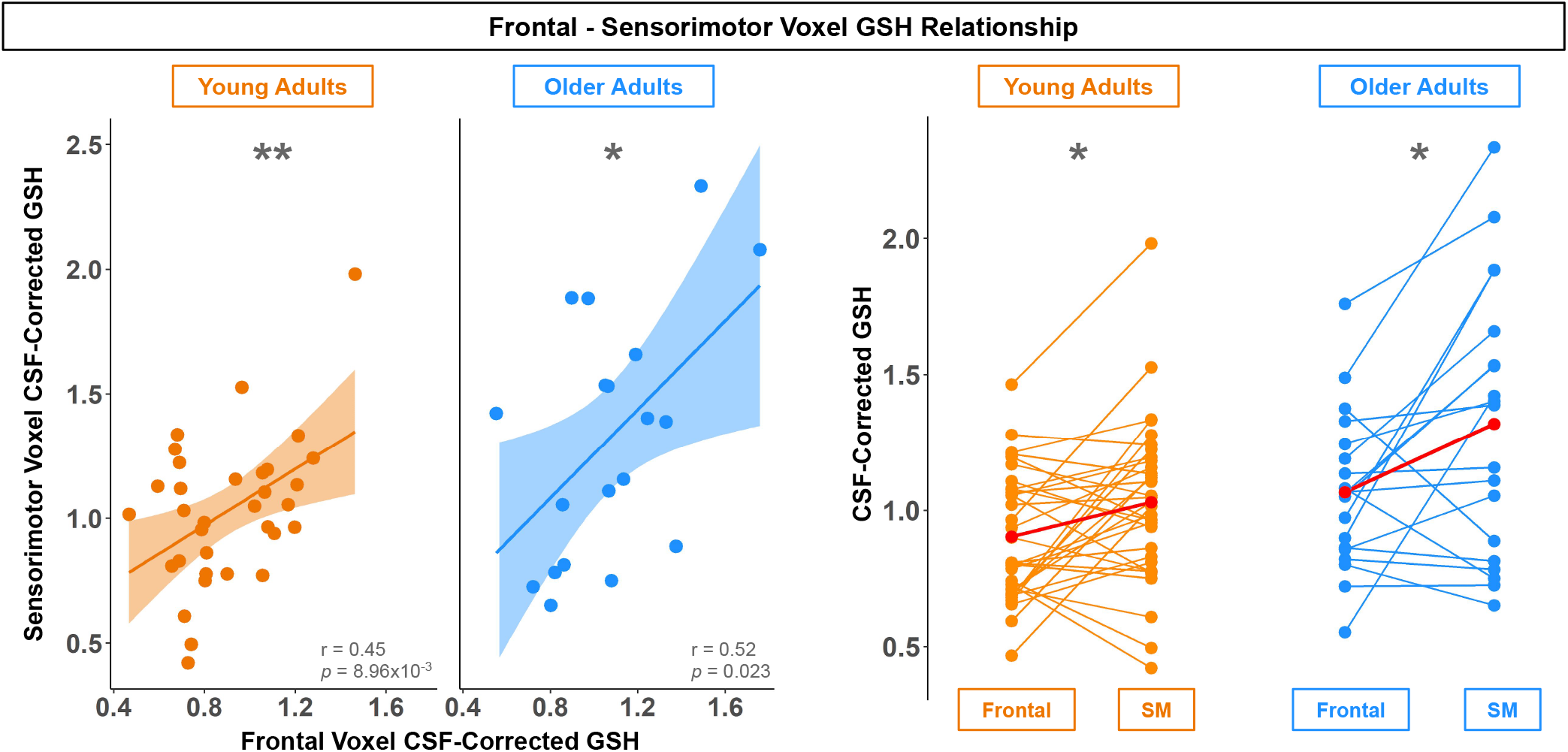
Higher GSH Levels in Sensorimotor versus Frontal Cortex. \**p* < 0.05; \*\**p* < 0.01. *Left*. Correlation of CSF-corrected GSH levels for the frontal and sensorimotor voxels for young (orange, left) and older (blue, right) adults. For both age groups, higher frontal GSH levels associated with higher sensorimotor GSH levels. *Right*. Frontal and SM (sensorimotor) voxel CSF-corrected GSH levels within each young (orange, left) and older (blue, right) adult. Each line represents one subject. For both age groups, GSH levels (group medians shown in red) were higher in the sensorimotor compared to the frontal voxel.

### GSH Relationships with Motor but Not Cognitive Performance for Older Adults Only

We did not observe relationships between frontal voxel GSH and performance, or between GSH and MoCA scores (Tables A4-A5). However, higher GSH levels within the sensorimotor, but not the frontal voxel, were associated with poorer performance on multiple motor measures for the older adults only.

Greater medial/lateral (M/L) sway speed and variability (i.e., greater postural instability) was correlated with higher GSH levels only for the older adults (Table A5; Fig. 4). The young adults had a weak but non-significant positive association between M/L sway speed and variability and GSH levels; there was a trend for an age difference in the partial correlation strength (*p* = 0.064).

**Fig. 4.**
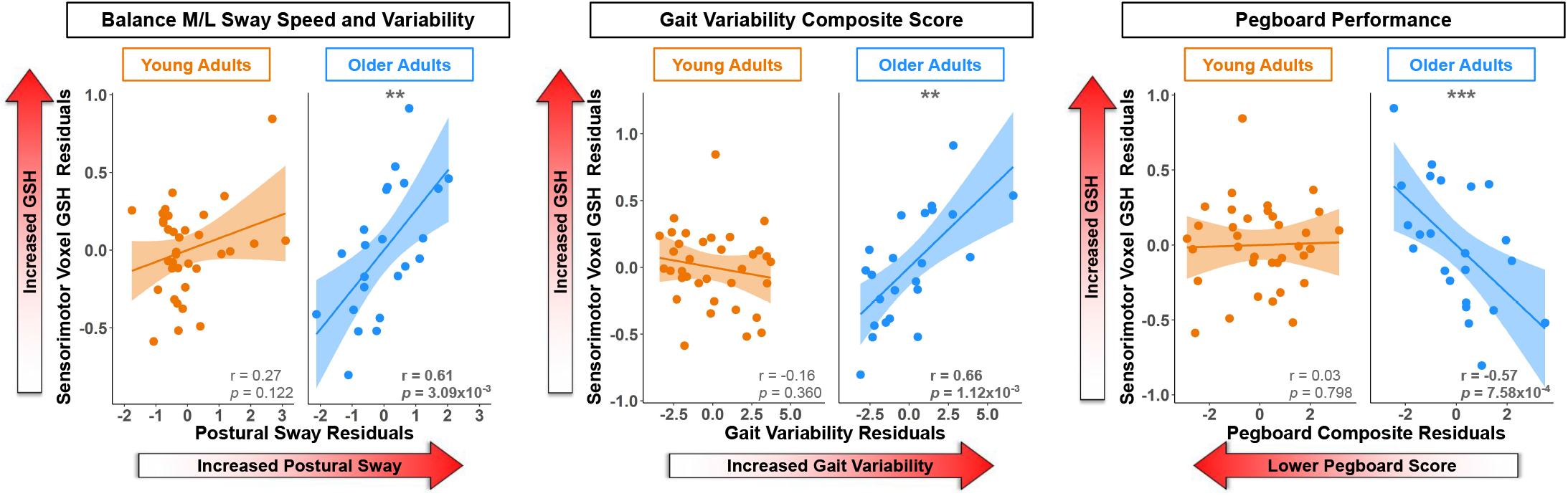
GSH Relationships with Motor but Not Cognitive Performance for Older Adults Only. \*\**p* < 0.01; \*\*\**p* < 0.001. Partial correlations of CSF-corrected GSH levels with M/L postural sway, gait variability, and pegboard performance for young (orange) and older (blue) adults. Partial correlations are accounting for the effects of the covariates included in each model. In each of these cases, there was a significant relationship between higher sensorimotor GSH levels and poorer motor performance for the older but not the younger adults.

Greater gait variability was correlated with higher sensorimotor GSH levels for the older adults only (Table A5; Fig. 4). No relationship emerged between gait variability and GSH levels for the young adults; the partial correlation strength was significantly different between young and older adults. As there is some evidence that walking speed contributes to gait variability (e.g., (Jordan et al., 2007)), we reran these models also including gait speed as a covariate; the relationship between sensorimotor GSH and gait variability for older adults remained significant (*p* = 0.004).

Poorer pegboard composite scores were associated with higher GSH levels only for the older adults (Table A5; Fig. 4), and there was a significant age difference in the partial correlation strength. Of note, although we measured sensorimotor GSH levels over the *lower limb* cortical representation, we still observed this GSH relationship with upper limb motor coordination. It was not the case that the pegboard composite score correlated with balance and gait (r = −0.04 and −0.08 for young adults; r = −0.23 and −0.31 for older adults; *p* > 0.05 in all cases); thus pegboard scores specifically index manual function.

To further test the specificity of the identified relationships between GSH and motor function for older adults, and not global shifts in metabolite concentrations, we reran the significant models above including as predictors the two other neurometabolites edited by HERMES: the excitatory neurochemicals glutamate + glutamine (Glx) and the primary inhibitory neurotransmitter within the brain, γ-aminobutyric acid (GABA). All relationships between physical function and GSH remained when including Glx and GABA as additional predictors; for older adults, the relationships remained significant between sensorimotor GSH levels and M/L sway speed/variability (*p* = 0.007), gait variability (*p* = 0.003), and manual dexterity (*p* = 0.008). There were no significant relationships between Glx or GABA levels and these motor metrics.

## Discussion

We identified higher CSF-corrected frontal and sensorimotor GSH levels for older compared to younger adults when accounting for age-related cortical atrophy. For both age groups, we identified a positive correlation between frontal and sensorimotor GSH levels, as well as higher GSH levels for the sensorimotor compared to the frontal voxel. For the older adults only, we identified multiple relationships between higher sensorimotor GSH levels and poorer motor performance.

One potential explanation for higher brain GSH levels for older adults is that higher levels of GSH occur as a compensatory response in an attempt to mitigate age-related increases in oxidative stress and maintain regional redox homeostasis within the brain. That is, perhaps in normal aging, in some regions of the brain, GSH antioxidant levels increase in response to increasing oxidative stress that occurs during aging. Past *in vivo* human studies have found higher MRS-measured GSH levels in MCI compared to age-matched controls (Duffy et al., 2014), but lower GSH levels in AD compared to controls (Mandal et al., 2015). Given the association between cognitive impairment and ROS production (Brawek et al., 2010), these findings could be interpreted as a ROS-induced compensatory upregulation of GSH in the early stages of cognitive decline. Similarly, past evidence suggests that MRS-measured GSH levels are higher in early schizophrenia (Wood et al., 2009), but lower after full symptoms emerge (Matsuzawa et al., 2008). Higher MRS-measured GSH levels have also been reported in post-traumatic stress disorder (Michels et al., 2014) and early psychosis (Godlewska et al., 2014). Further, pharmacologically-induced GSH depletion in the brain has been shown to result in cognitive decline in rodents (González-Fraguela et al., 2018). Therefore, high GSH levels could be associated with high levels of underlying cellular stress (e.g., ROS emissions) until reaching a threshold that exceeds the hormetic response capabilities of the cell.

The precise mechanisms that dictate GSH regulation in the aging brain remain unknown. Nonetheless, in theory, an increase in GSH could reflect either upregulation of GSH production or downregulation of GSH catabolism. For instance, one study (Mythri et al., 2011) found significantly less γ-glutamyl transpeptidase (γ-GT) activity in the cortex and striatum of Parkinson’s disease patients, suggesting that lower rates of GSH breakdown were contributing to the increased GSH seen in these tissue samples.

Increased oxidative stress with aging could also explain increases in GSH. Increases in cellular antioxidants in response to oxidative insults have been well documented. Multiple animal model and cell culture studies have shown a compensatory upregulation of GSH in response to oxidative stress (Ong et al., 2000), including exposure to toxins such as mercury (Hoffman et al., 2005), radiation (Di Toro et al., 2007), neonatal alcohol (Smith et al., 2005), and methamphetamine (Harold et al., 2000), as well as neurological diseases such as models of Parkinson’s disease (Aluf et al., 2010; Rodríguez Navarro et al., 2007), Huntington’s disease (Tkac et al., 2007), and AD (Tchantchou et al., 2005).

While aberrant ROS production can be detrimental to cellular health, ROS also serve as signaling molecules capable of regulating transcriptional events in the cell. The Keap1/Nrf2 signaling pathway is a canonical oxidative stress sensor in the cell. Nrf2 is a key protein responsible for increasing antioxidant enzymes by regulating transcription of antioxidant-related genes (Itoh et al., 1999). The endogenous protein, Keap1, suppresses Nrf2 activity under basal conditions by facilitating its removal from the cell. However, oxidative modification of Keap1 by ROS removes its inhibitory effects on Nrf2 and allows for Nrf2-mediated transcription of antioxidant related genes (Sekhar et al., 2010). Importantly, Nrf2 regulates transcription of the rate-limiting enzyme responsible for synthesizing GSH, γ-glutamate-cysteine ligase (Yang et al., 2005). Therefore, increased ROS emissions with brain aging may result in increased GSH via the Keap1/Nrf2 signaling pathway. There is some literature support for this idea; for instance, exposure of astrocytes in cell culture to high levels of ROS induces transcription of genes responsible for synthesizing GSH (Gegg et al., 2003; Sagara et al., 1996).

It could also be that the observed GSH changes relate to changes in cell type abundance within the aging brain. As GSH is present in higher concentrations in glia compared to neurons (Rice & Russo-Menna, 1997), increasing GSH levels could be associated with the increased gliosis that occurs with brain aging (Tong et al., 2011). Stereological cell counting in post-mortem human brain suggests that the abundance of astrocytes, one of the predominant producers of brain GSH, remains constant throughout aging while the abundance of other cell types (e.g., oligodendrocytes) decreases (Pelvig et al., 2008). This may be particularly important given that astrocytes are also more capable of inducing the antioxidant defense response via Nrf2 signaling compared to other neuronal cell types (Baxter & Hardingham, 2016).

This finding of higher CSF-corrected GSH levels for older compared to younger adults is in line with the results of Tong and colleagues (Tong et al., 2016). This group identified GSH increases across the lifespan (i.e., 1 day to 99 years old) in post-mortem frontal cortex. However, this finding is in contrast to the results of Emir and colleagues (2011) who reported lower occipital cortex GSH levels for older compared to younger adults using edited MRS at 4T. There are several key differences between our work and this study. Emir and colleagues examined a different brain region, and their elderly sample (76.6 ± 6.1 years) was older than ours; these factors likely contributed to their differing results. More recent work by this group using non-edited MRS at 7T found no age differences in posterior cingulate or occipital cortex GSH levels (Marjańska et al., 2017); however, again, this study tested different brain regions and an older sample compared to our work. The lack of GSH age differences in posterior brain regions (but not frontal or sensorimotor cortex) could also be due in part to the well-established posterior to anterior shift of brain activity with aging (Davis et al., 2008; Jockwitz et al., 2019). It could be that reduced neural signaling within posterior brain areas leads to lower regional GSH levels; that is, compensatory upregulation of GSH has ceased in these posterior regions as the hormetic response capabilities of these cells have been exceeded.

Given the limited age range in the present study, it is unknown whether the apparent age-related increase in brain GSH presented here may abate in extreme conditions of oxidative stress, such as neurological disease or very old age, as the compensatory response is overwhelmed (e.g., as recycling or *de novo* synthesis mechanisms are compromised). Future longitudinal studies and enrollment of much older adults would clarify this.

Some work has reported regional differences in cortical GSH levels (Nezhad et al., 2017; Srinivasan et al., 2010; Tong et al., 2016). Here we found a positive correlation between frontal and sensorimotor CSF-corrected GSH levels for both young and older adults, as well as higher GSH levels in the sensorimotor compared to the frontal voxel for both age groups. However, we identified tissue composition differences between the two voxels for the younger adults only. Young adults had higher gray matter and CSF and lower white matter fractions in the sensorimotor compared to the frontal voxel. For older adults, there were no tissue composition differences between voxels. These young adult findings fit with one past study reporting higher GSH concentrations in voxels with more gray matter than white matter (Srinivasan et al., 2010). However, another study (Nezhad et al., 2017) reported conflicting findings of higher GSH concentrations in the cortical region with less gray matter (i.e., anterior cingulate versus occipital cortex). Thus, as we did not find voxel composition differences for the older adults, but we did find higher sensorimotor versus frontal GSH levels, we suspect that (as discussed by Rae and Williams (2017)), GSH levels likely vary across brain region, but in a more complex manner than that which reflects only gray and white matter differences. This notion is further supported by recent work suggesting that human primary motor and somatosensory cortices show proportionally steeper trajectories of volume, myelin, and iron declines with advancing age compared to other brain regions (Taubert et al., 2020). It could be that the sensorimotor cortex structure and neurochemical composition is affected more or earlier by oxidative stress compared to other brain regions.

There were no associations between GSH levels and cognitive performance (i.e., MoCA scores). Our past work (Porges, Woods, Edden, et al., 2017) suggests that MoCA scores are sensitive enough to identify associations between MRS-measured neurometabolites and cognitive status. In contrast to our previous work (*n* = 93 older adults; mean age = 73.2 ± 9.9), here we included fewer participants, although our older adult ages were similar. Additionally, participants in the present sample had higher MoCA scores compared to our previous work (mean = 25.5 ± 2.5). It could be that, among this higher-functioning older adult cohort, we did not have enough variation in MoCA scores to identify a significant association. Furthermore, the limited past work in normal aging has failed to find any relationships between GSH levels and cognitive status (Chiang et al., 2017; Emir et al., 2011); such a relationship has previously been identified only in pathological conditions such as MCI and AD (Mandal et al., 2015; Mandal et al., 2012; Oeltzschner et al., 2019). Thus, it could be that GSH-cognition relationships only emerge in cases of more severe cognitive decline, when brain resources (such as antioxidant availability) have substantially declined.

We found several associations between higher GSH levels and poorer motor performance (i.e., greater M/L postural sway, greater gait variability, and poorer manual dexterity). These motor performance variables have functional significance. Greater M/L postural sway (Maki et al., 1994; Stel et al., 2003) and greater gait variability (e.g., (Hausdorff et al., 2001)) associate with a greater risk of falling for older adults. Declines in manual function are associated with decreased independence for older adults (Falconer et al., 1991; Williams et al., 1982). Although relationships of GSH levels with motor performance in normal aging have not been previously investigated, these findings fit with past work that identified relationships between GSH levels and movement disorders (Choi et al., 2015; Doss et al., 2015; Srinivasan et al., 2010; Weerasekera et al., 2019; Weiduschat et al., 2014). Together, these findings may indicate that GSH is providing a compensatory response to increasing oxidative stress and related tissue damage in the normally aging brain. Higher sensorimotor versus frontal cortex GSH levels (discussed above) could suggest that the sensorimotor cortex is disproportionately affected by oxidative stress in older age (and thus requires the largest GSH antioxidant response). This regionally heightened oxidative stress may then be contributing to these age-related declines in motor function.

Importantly, we identified relationships between motor performance and sensorimotor but not frontal GSH levels. This suggests regional specificity for these GSH relationships, rather than poorer motor performance being a consequence of increased oxidative stress throughout the brain. Moreover, we found no relationship between cortical Glx or GABA levels and these motor performance metrics, again supporting the specificity of this GSH relationship with motor behavior for older adults and not result of more general age related shifts in metabolite concentrations.

Although we measured sensorimotor GSH levels over the lower limb cortical representation, we still identified a GSH relationship with upper limb motor coordination. The pegboard composite score did not correlate with gait or balance performance, suggesting that it represents a unique motor measure, which independently associates with sensorimotor GSH. It could be that lower limb sensorimotor GSH levels are related to upper limb sensorimotor GSH levels; this is probable given that we did not find any associations between motor performance and frontal GSH levels, and that we found positive correlations between GSH levels across the two voxels. However, this remains to be examined in future studies.

There are several limitations to the present work. We included fewer older adults than originally anticipated due to the COVID-19 global pandemic; however, based on our power analysis, we were still adequately powered to test age group differences in GSH. In addition, our cross-sectional approach precluded us from assessing how GSH levels alter with aging or how changes in GSH levels across the lifespan relate to declines in motor performance. There is some evidence that diet may influence MRS-measured GSH levels. One study (Choi et al., 2015) found an association between dairy consumption and brain GSH levels among older adults. In the present work, we did not record food intake or restrict diet prior to the MRI scan; future studies should characterize any effects of diet on brain GSH levels. Finally, other general limitations of MRS, such as the large voxel size required, are currently unavoidable with this methodology.

These results provide insight into the association between brain aging and oxidative stress. We demonstrate higher CSF-corrected GSH levels with normal aging, suggesting a GSH compensatory response to increased oxidative stress with older age. We report higher GSH levels in the sensorimotor cortex compared to the frontal cortex for both age groups, as well as multiple associations between sensorimotor CSF-corrected GSH levels (but not GABA or Glx levels) and poorer balance, gait, and manual dexterity. Together, these results suggest that MRS-measured GSH could be a marker of neural compensation for increased oxidative stress with brain aging and also a marker of poorer motor performance. These results could stem from greater or earlier effects of oxidative stress on the sensorimotor compared to the frontal cortex.

## Materials and Methods

The University of Florida’s Institutional Review Board provided ethical approval for the study, and all participants provided their written informed consent at the first testing session.

### Participants

We recruited 37 young and 23 older adults from the Gainesville, FL community. Exclusion criteria included: history of any neurologic condition (e.g., stroke, Parkinson’s disease, seizures, or a concussion in the last six months) or psychiatric condition (e.g., active depression or bipolar disorder). We also excluded those who self-reported smoking, consuming more than two alcoholic drinks per day on average or a history of treatment for alcoholism. All subjects were screened for magnetic resonance imaging (MRI) eligibility; we excluded those with any contraindications (e.g., implanted metal, claustrophobia, or pregnancy). All subjects were right-handed and self-reported an ability to walk unassisted for at least 10 minutes and to stand for at least 30 seconds with their eyes closed. Participants disclosed all current prescribed and over-the-counter medications.

Prior to enrollment, we screened participants for suspected cognitive impairment over the phone using the Telephone Interview for Cognitive Status (TICS-M) (de Jager et al., 2003). We excluded those who scored <21 of 39 points; this is equivalent to scoring <25 points on the Mini-Mental State Exam (MMSE) and indicates probable cognitive impairment (de Jager et al., 2003). At the first testing session, participants were re-screened for cognitive impairment using the Montreal Cognitive Assessment (MoCA) (Nasreddine et al., 2005); we excluded those who scored <23 of 30 points (Carson et al., 2018).

### Sample Size

Due to the COVID-19 global pandemic, data collection was terminated before we completed the recruitment of older adult subjects. However, based on a power analysis, 37 young and 23 older adults is more than sufficient for detecting an age difference in MRS-measured GSH levels. We calculated the minimum necessary sample size using G*Power 3.1 (Erdfelder et al., 1996). We based this calculation on the only past study testing age differences in MRS-measured GSH (Emir et al., 2011); this study reported an effect size of *d* = 1.65 for age differences in occipital cortex GSH levels (Emir et al., 2011). With power = 0.80 and ⍺ = 0.05, a two-sample independent *t*-test (i.e., to characterize group age differences in GSH levels) would require only six subjects per group.

### Testing Sessions

Prior to the first session, we collected basic demographic information, including age, sex, years of education, and medical history, as well as information regarding self-reported exercise, handedness, and footedness. We also collected basic anthropometric information, such as height, weight, and leg length.

Participants then completed behavioral testing, followed by an MRI session approximately one week later (Fig. 5). For 24 hours prior to each session, participants refrained from consuming alcohol, nicotine, or any drugs other than the medications they previously disclosed. At the start of each session, participants completed the Stanford Sleepiness Questionnaire, which asks for the number of hours slept the previous night and for a rating of current sleepiness (Hoddes et al., 1972).

**Fig. 5.**
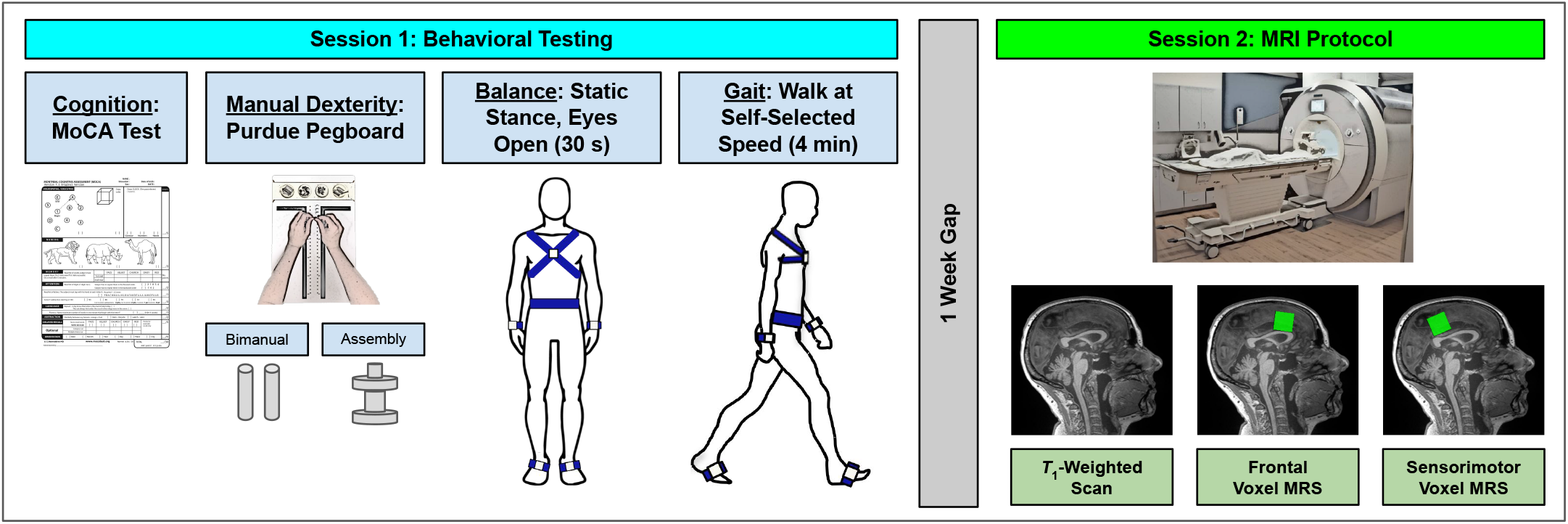
Methods Overview. <u>Left</u>. During Session 1, participants first completed the MoCA test of cognition. Participants then completed two manual dexterity tasks: the Purdue pegboard bimanual and assembly conditions. Next, participants were instrumented with six IMUs (sensors are pictured in gray with blue straps) and completed a 30-s balance task in which they stood as still as possible with their eyes open, gazing at a blank white wall. Subjects then completed a 4-minute walk at a self-selected speed across a 32-foot room. <u>Right</u>. During Session 2, participants completed an MRI protocol which included a *T*_1_-weighted anatomical scan and two edited MRS scans to quantify neurometabolites in a frontal and sensorimotor voxel.

#### Session 1: Behavioral Testing

##### MoCA Test

Participants first completed the MoCA (Nasreddine et al., 2005). We added one point to the scores of participants with ≤12 years of education (Nasreddine et al., 2005).

##### Balance Task

Participants completed the four-part Modified Clinical Test of Sensory Interaction in Balance (m-CTSIB) while instrumented with six Opal inertial measurement units (IMUs; v2; APDM Wearable Technologies Inc., Portland, OR, USA). IMUs were placed on the feet, wrists, around the waist at the level of the lumbar spine, and across the torso at the level of the sternal angle (Fig. 5). Participants stood as still as possible facing a blank white wall for four 30-second trials: 1) eyes open; 2) eyes closed; 3) eyes open, foam surface; and 4) eyes closed, foam surface. Here we report only on performance during the eyes open condition. We elected to use only the eyes-open condition because several subjects had scores greater than ±3 standard deviations from the group mean for the foam conditions; thus, using the eyes-open condition prevented us from needing to exclude any outlier data. Furthermore, previous work has reported age differences in postural sway during quiet stance with eyes open (Baloh et al., 1994; Maki et al., 1990), and eyes open postural sway has been shown to predict falls among older adults (Fernie et al., 1982; Maki et al., 1990).

Inertial data were recorded using MobilityLab software (version 2; APDM Wearable Technologies Inc., Portland, OR, USA). After each trial, MobilityLab calculated 25 spatiotemporal features of postural sway (Table B1) using the validated iSway algorithm (Mancini et al., 2012). To condense these variables into several summary metrics, we ran an exploratory factor analysis (Appendix B). This procedure yielded four factors: anterior/posterior (A/P) sway path, A/P sway speed and variability, M/L sway path, and M/L sway speed and variability. We then calculated a balance composite score for each factor to use in subsequent analyses.

##### Four-Minute Walk

While instrumented with the IMUs, participants also completed an overground walk. Participants walked back and forth across a 32-foot room for four minutes at whichever pace they considered to be their “normal walking speed.” Participants were instructed to refrain from talking, to keep their arms swinging freely at their sides, and to keep their head up and gaze straight ahead. Each time they reached the end of the room, they completed a 180-degree turn and walked the length of the room again.

After the session, the MobilityLab software calculated 14 spatiotemporal gait variables of interest (Table C1). The algorithm for calculating these metrics has been validated through comparison to force plate and motion capture data (see internal validation by MobilityLab: https://support.apdm.com/hc/en-us/articles/360000177066-How-are-Mobility-Lab-s-algorithms-validated- and (Washabaugh et al., 2017)). To obtain summary metrics of gait, we extracted one variable from each of the four gait domains described by Hollman and colleagues (Hollman et al., 2011): gait rhythm (cadence (steps/min)), gait phase (stance (% gait cycle)), gait pace (composite score), and gait variability (composite score). See Appendix B for further details regarding the selection and calculation of these summary metrics.

##### Pegboard Tasks

Participants completed two tasks using a Purdue pegboard (Lafayette Instruments, Lafayette, IN, USA). For the bimanual task, participants had 30 seconds to place as many pegs as possible into the slots; in this case, participants used both hands at the same time to place peg pairs (Fig. 5). Scores were based on the number of completed peg pairs. For the assembly task, participants had one minute to complete as many “assemblies” as possible (Fig. 5). An assembly consisted of using both hands to piece together metal pins, collars, and washers. Scores were based on the number of completed assemblies. These tasks were selected as they each require complex coordination of both hands and performance declines with age (Agnew et al., 1988; Vasylenko et al., 2018). For further analysis, we created a composite score of pegboard performance by converting the bimanual and assembly scores to standardized *Z-* scores and then taking the sum of these two *Z-*scores.

### Session 2: MRI Scan

#### T1 Acquisition

MRI was conducted using a Siemens MAGNETOM Prisma 3T scanner (Siemens Healthcare, Erlangen, Germany) using a 64-channel head coil. We first collected a 3D *T*_1_-weighted anatomical image using a magnetization-prepared rapid gradient-echo (MPRAGE) sequence for MRS voxel placement and tissue segmentation/correction. The parameters for this anatomical image were as follows: TR = 2000 ms, TE = 3.06 ms, flip angle = 8°, FOV = 256 × 256 mm^2^, slice thickness = 0.8 mm, 208 slices, voxel size = 0.8 mm^3^.

#### MRS Acquisition

In the following sections and in Table A2, we describe all parameters suggested by the Magnetic Resonance Spectroscopy quality assessment tool (MRS-Q) (Peek et al., 2020). We used the universal Hadamard Encoding and Reconstruction of MEGA-Edited Spectroscopy (HERMES) sequence to simultaneously detect GSH, GABA, and Glx (Saleh et al., 2016; Saleh et al., 2019). HERMES is a *J*-difference editing method that allows for multiple MEGA-PRESS (Mescher et al., 1998) experiments to be conducted simultaneously. Briefly, the HERMES sequence includes four sub-experiments containing: A) a dual-lobe editing pulse, ON_GABA_ = 1.9 ppm, ON_GSH_ = 4.56 ppm, and three single-lobe editing pulses: B) ON_GABA_ = 1.9 ppm, C) ON_GSH_ = 4.56 ppm, and D) OFF_GABA_, OFF_GSH_ = 7.5 ppm. The Hadamard combination A-B+C-D derives GSH-edited spectra, and A+B-C-D derives GABA+- and Glx-edited spectra. Additional HERMES parameters included: total acquisition time = 10:48 minutes, TR = 2000 ms, TE = 80 ms, 20-ms editing pulse duration, averages = 320, 2048 data points, 2 kHz spectral width, and variable power and optimized relaxation delays (VAPOR) water suppression. Shimming was performed using the Siemens interactive shim tool and FAST(EST)MAP (Gruetter, 1993).

We collected data from two 30 × 30 × 30 mm^3^ voxels in the medial frontal cortex and bilateral sensorimotor cortex (Fig. 6). We placed the frontal voxel superior to the genu of the corpus callosum on the mid-sagittal slice. We placed the sensorimotor voxel to align with the lower limb primary sensorimotor cortex. We aligned the center of this voxel with the posterior portion of the motor hand knob in the axial view, then centered the voxel on the midline of the brain, and placed the voxel as superior as possible while still remaining on brain tissue.

**Fig. 6.**
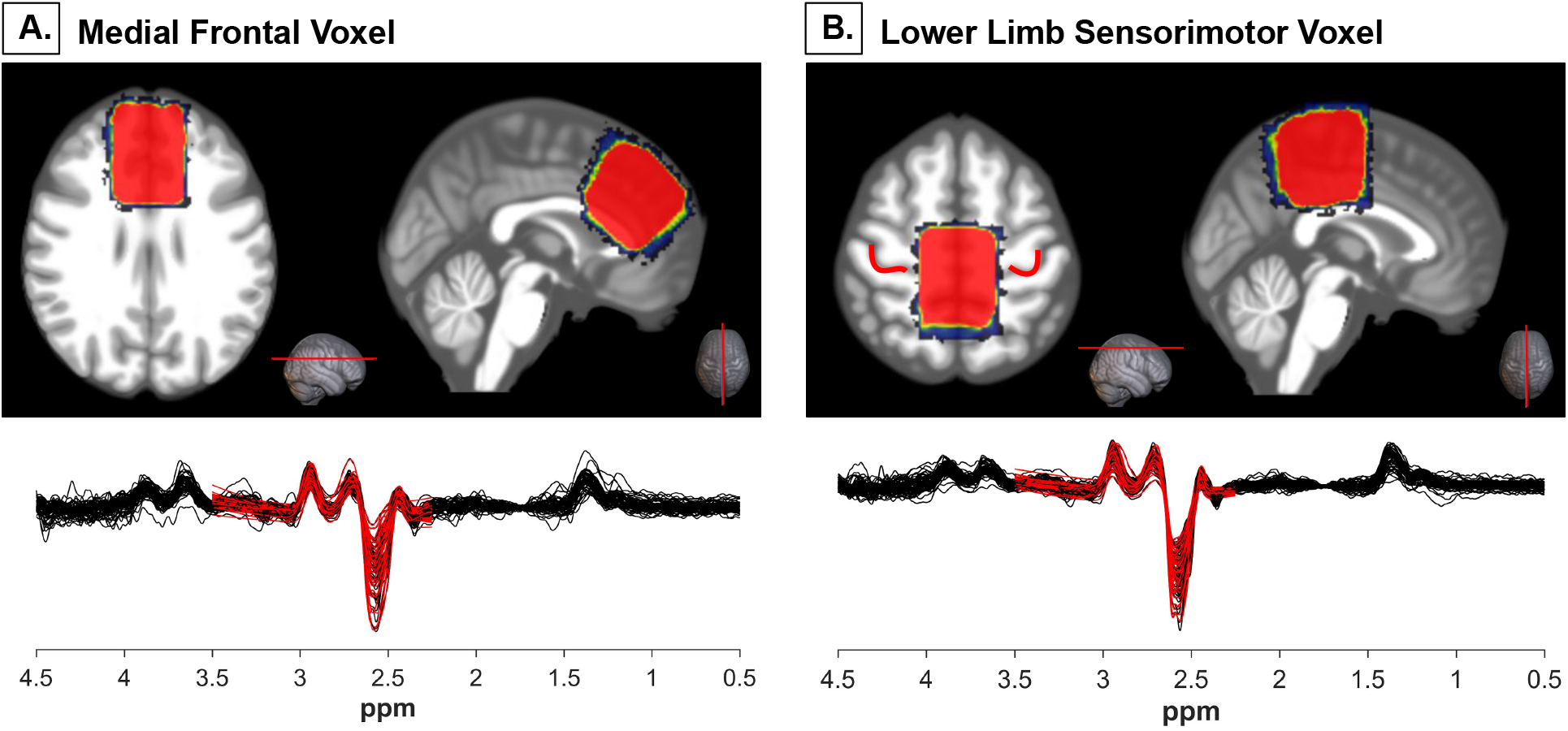
MRS Voxel Placement. *A*. <u>Top</u>. Placement of the medial frontal voxel: superior to the genu of the corpus callosum on the midsagittal line. The voxel shown is every subject’s voxel, normalized to standard space and overlaid onto a template brain. Warmer colors indicate areas of more overlap across subjects. <u>Bottom</u>. Plot created using `PaperPlot.m` showing all participants’ spectra overlaid (black) and the GSH model fit (red) for the frontal voxel. *B*. <u>Top</u>. Placement of the sensorimotor voxel: centered with the posterior portion of the motor hand knobs in the axial view (motor hand knobs are outlined in red), then centered on the midline of the brain and placed as superior as possible. The voxel shown is every subject’s voxel, normalized to standard space and overlaid onto a template brain. Warmer colors indicate areas of more overlap across subjects. <u>Bottom</u>. Plot created using `PaperPlot.m` showing all participants’ spectra overlaid (black) and the GSH model fit (red) for the sensorimotor voxel.

#### MRS Processing

We analyzed MRS data using Gannet (version 3.1.5) (Edden et al., 2014) in MATLAB (R2019b). First, we ran the *GannetLoad.m* and *GannetFit.m* functions, which include: 1) coil combination using generalized least squares (An et al., 2013); 2) estimation of the *B*_0_ drift using the creatine (Cr) signal at 3 ppm; 3) robust spectral registration to minimize subtraction artifacts (Mikkelsen et al., 2018); 4) Hadamard-combination of the fully processed HERMES sub-spectra to generate GSH- and GABA+-edited difference spectra; 5) application of the Hankel singular decomposition water filtering method to remove the residual water signal (Barkhuijsen et al., 1987); and 6) implementation of a weighted nonlinear regression to model the two difference-edited signals; here, the neighboring co-edited signals were downweighted to reduce their impact on modeling errors. The GSH-edited spectrum was modeled between 2.25 and 3.5 ppm using a Gaussian to model the GSH signal at 2.95 ppm, four Gaussians to model the coedited aspartyl signals at 2.55 ppm, and a nonlinear baseline.

We used *GannetCoRegister.m* to create a binary mask of the MRS voxels and register these masks to the *T*_1_-weighted structural image. We then used the Computational Anatomy Toolbox 12 (CAT12, version 1450) (Gaser & Dahnke, 2016) to segment each subject’s *T*_1_-weighted image. We implemented *GannetSegment.m*, which uses segmentation results to determine voxel tissue fractions (i.e., fractions of gray matter, white matter, and CSF) and to correct GSH estimates for tissue composition (Harris et al., 2015). Correcting for tissue composition enhances the interpretation of MRS data. Metabolite levels, as well as reference signals, differ between gray matter, white matter, and CSF (Harris et al., 2015). Tissue correction is particularly relevant for aging populations (Porges, Woods, Lamb, et al., 2017). For instance, if older adults have less gray matter due to age-related atrophy in a voxel compared to young adults, the older adults will also present with less metabolite concentration in that voxel. Correcting for tissue composition thus permits assessment of whether there are age differences in neurometabolite levels in the tissue that remains in the voxel. Throughout the present work, we report CSF-corrected GSH levels referenced to water.

#### MRS Exclusions

See Table E1 for details on exclusions of MRS datasets. We excluded MRS datasets if the GSH fit error (i.e., `GSH.FitError_W`) was greater than 20% or if robust spectral registration failed for that dataset. We selected 20% for several reasons: 1) datasets with fit errors <20% passed acceptable visual inspection and 2) fit errors ≥20% were >2.5 standard deviations above the group mean (i.e., >97th percentile). Thus, similar to (Saleh et al., 2020), we selected a threshold value for data rejection. Of note, we did not exclude one older adult for whom we used a 20-channel head coil instead of a 64-channel coil due to his large head size. The uncorrected and CSF-corrected GSH levels for this individual fell within the range of that of the other older subjects. See Fig. C1 for details.

### Statistical Analyses

We conducted all statistical analyses using R (version 4.0.0) (R Core Team, 2013).

#### Age Group Comparisons

For each analysis involving comparisons between the age groups, we first tested the parametric *t*-test assumptions of normality within each group (using `shapiro.test`) and homogeneity of variances between the groups (using `leveneTest` in the `car` package (Fox & Weisberg, 2018)). We then tested age group differences, as described below.

##### Parametric Tests

The majority of MRS variables met the required assumptions, so we used `t.test` to conduct parametric, independent-samples, two-sided *t*-tests. For each MRS variable, we report *t*-test results, in addition to group means, standard deviations, and Cohen’s *d* as a measure of effect size.

##### Nonparametric Tests

In several cases (i.e., age differences in demographic information and cognitive/motor performance), the majority of variables did not meet parametric *t*-test assumptions, so we instead used `wilcox.test` to conduct nonparametric, independent-samples, two-sided Wilcoxon rank-sum tests for group differences. In these cases, we report the group medians and interquartile ranges for each demographic variable. We also report nonparametric effect sizes (Field et al., 2012; Rosenthal et al., 1994); see Appendix F for details on this calculation. To test for differences in the sex distribution within each age group, we conducted a Pearson chi-square test using `chisq.test`.

#### Within-Subject Tests

To examine within-subject differences in MRS variables between the frontal and sensorimotor voxels, within each age group, we implemented parametric, paired-samples *t*-tests. These variables met the normality assumption required for parametric paired *t*-tests

#### GSH Correlations

We conducted Pearson correlations using `cor.test` to assess the relationship between frontal and sensorimotor GSH levels. These metrics met the assumptions of linear covariation and normality. Here we also performed a Fisher *r*-to-*Z* transformation on the correlation coefficient and then tested for a difference in correlation strength between the age groups using a one-sided `r.test` in the `psych` package (Revelle, 2014).

#### GSH Relationships with Cognitive and Motor Performance

For each behavioral metric, we used `lm` to test the relationship between CSF-corrected GSH levels and performance for both voxels and age groups. For the MoCA score models, we controlled for sex and years of education (Malek-Ahmadi et al., 2015). For the balance and gait models, we controlled for sex and leg length, as these each affect postural sway (e.g., (Kim et al., 2010)) and gait (e.g., (Ko et al., 2011; Kobayashi et al., 2016; Samson et al., 2001)). For the manual dexterity model, we controlled for sex, as there is some evidence of sex differences in Purdue pegboard performance (Vasylenko et al., 2018).

We also computed the partial correlation for each GSH-performance relationship (i.e., the correlation controlling for the covariates listed above) by correlating the residuals from 1) regressing each of the covariates (but not GSH concentration) onto the performance variable, and 2) regressing each of the covariates onto GSH concentration. Finally, as described above, we used a Fisher *r*-to-*Z* transformation to test for age differences in the strength of the partial correlation. As several variables did not meet the linear regression assumptions of heteroscedasticity and normality, for each model that yielded a significant GSH-performance relationship, we also ran a nonparametric version of that model using `npreg` and `npsigtest` in the `np` package (Hayfield & Racine, 2008).

As noted in the Results section, we reran the gait variability model including gait speed as an additional covariate because there is some evidence that walking speed contributes to gait variability (e.g., (Jordan et al., 2007)). Further, for each model that indicated a significant relationship between GSH levels and behavior, we reran the model also including GABA and Glx as covariates. This was to provide further support for the specificity of the relationship between GSH levels and motor performance; that is, we hypothesized that these neurometabolites would not relate to behavior, and that including these would not influence the significant relationship between GSH levels and motor performance.

#### Corrections for Multiple Comparisons

We corrected *p-*values within each results table using `p.adjust` with `method = “bh”` to apply the Benjamini-Hochberg FDR correction (Benjamini & Hochberg, 1995). We present the uncorrected *p*-values within the tables and describe in the table footnote if any *p*-values did not pass the FDR correction. With the exception of two cases in which the *p*-values were *p* < 0.10, all other results remained significant (*p* < 0.05) after applying the FDR correction for multiple comparisons.

## Acknowledgements

During completion of this work, KH was supported by a National Science Foundation Graduate Research Fellowship under Grant no. DGE-1315138 and DGE-1842473 and by NINDS training grant T32-NS082128. EP was supported by NIAAA grant K01 AA025306 and the McKnight Brain Research Foundation; the Center for Cognitive Aging and Memory at the University of Florida. A portion of this work was performed in the McKnight Brain Institute at the National High Magnetic Field Laboratory’s Advanced Magnetic Resonance Imaging and Spectroscopy (AMRIS) Facility, which is supported by National Science Foundation Cooperative Agreement No. DMR-1644779 and the State of Florida. This work was also supported in part by an NIH award, S10OD021726, for High End Instrumentation. This study applied tools developed under NIH grants R01 EB016089, R01 EB023963 and P41 EB015909.

The authors wish to thank Aakash Anandjiwala, Justin Geraghty, and Alexis Jennings-Coulibaly for their help in subject recruitment and data collection. The authors also wish to thank all of the participants who volunteered their time, as well as the McKnight Brain Institute MRI technologists, without whom this project would not have been possible.

## Author Contributions

KH participated in initial study design, collected all data, processed the MRS data, conducted the statistical analyses, created the figures, and wrote the manuscript. HH contributed to manuscript writing and results interpretation. PAJ assisted with data collection, data processing, and manuscript preparation. MM advised on MRS data processing and methods, in addition to contributing to manuscript preparation. CH consulted on the design and analysis of the motor performance tests. RE advised on MRS data acquisition, processing, interpretation, and manuscript preparation. RS and EP oversaw study design and led the interpretation and discussion of results. All authors participated in revision of the manuscript.

