## Supplementary Materials for "*In Vivo* Brain Glutathione is Higher in Older Age and Correlates with Mobility"

### **Appendix A:** Supplemental Tables

**Table A1**. Participant Demographics, Physical Characteristics, Sleep, and Testing Timeline

| | **Young Adults** | **Older Adults** | $W$ **or** $\chi^2$ | $p$ | **Nonparametric Effect Size**[a] |
|---|---|---|---|---|---|
| **Demographics** | | | | | |
| Sample Size | 37[b] | 23[b] | -- | -- | -- |
| Age | 21.8 (2.5) | 72.8 (8.9) | 0.00 | **$1.04 \times 10^{-10}$ ***** | **-0.83** |
| Sex | 19 F; 18 M | 11 F; 12 M | 0.07 | 0.791 | -- |
| Years of Education | 15.0 (3.0) | 16.0 (3.0) | 219.0 | **$1.53 \times 10^{-3}$ **** | **-0.41** |
| Alcohol Use[c] | 2 (3.0) | 2 (4.0) | 480.0 | 0.406 | -0.11 |
| **Physical Characteristics and Fitness** | | | | | |
| Handedness Laterality Score[d] | 85.7 (25.0) | 100.0 (22.4) | 351.0 | 0.240 | -0.15 |
| Footedness Laterality Score[d] | 100.0 (22.2) | 100.0 (133.9) | 492.5 | 0.268 | -0.14 |
| Body Mass Index (BMI) | 22.7 (5.6) | 26.0 (3.9) | 184.5 | **$2.55 \times 10^{-4}$ ***** | **-0.47** |
| Leisure-Time Physical Activity[e] | 46.0 (38.0) | 29.0 (21.0) | 563.5 | **$1.44 \times 10^{-2}$ *** | **-0.32** |
| **Sleep** | | | | | |
| Hours of Sleep, Behavioral Session | 7.0 (1.5) | 7.5 (1.5) | 385.0 | 0.537 | -0.08 |
| Sleepiness Rating[†], Behavioral Session | 2.0 (1.0) | 1.0 (1.0) | 594.0 | **$5.66 \times 10^{-3}$ **** | **-0.36** |
| Hours of Sleep, MRI Session | 7.0 (2.0) | 7.0 (1.5) | 324.5 | 0.123 | -0.20 |
| Sleepiness Rating[†], MRI Session | 2.0 (1.3) | 1.0 (1.0) | 585.5 | **$4.39 \times 10^{-3}$ **** | **-0.37** |
| **Testing Timeline** | | | | | |
| Time Between Behavioral Testing and MRI (# of Days) | 4.0 (7.0) | 4.0 (4.5) | 415.0 | 0.878 | -0.02 |
| Difference in Start Time for Behavioral Testing vs. MRI (hours) | 1.3 (1.5) | 1.2 (1.1) | 462.5 | 0.579 | -0.07 |

*Note.* *$p < 0.05$, **$p < 0.01$, ***$p < 0.001$. In the second and third columns, we report the median ± interquartile range (IQR) for each age group in all cases except for sex. For sex, we report the number of males and females in each age group. In the fourth and fifth columns, for all variables except sex, we report the result of a nonparametric two-sample, two-sided Wilcoxon rank-sum test. For sex, we report the result of a Pearson's chi-square test for differences in the sex distribution within each age group. All significant $p$-values (bolded) remained significant at $p < 0.05$ after applying the Benjamini-Hochberg FDR correction (Benjamini & Hochberg, 1995).

[a] In the sixth column, we report the nonparametric effect size as described by (Field et al., 2012; Rosenthal et al., 1994); see Appendix F.

[b] All subjects (i.e., 37 young and 23 older adults) are included in the comparisons in this table. However, we excluded several individuals from the subsequent analyses involving the MRS and behavioral metrics. See Table E1 for details.

[c] Participants self-reported alcohol use on the Alcohol Use Disorders Identification Test (AUDIT) (Piccinelli, 1998).

[d] We calculated handedness and footedness laterality scores using the Edinburgh Handedness Inventory (Oldfield, 1971) and Waterloo Footedness Questionnaire (Elias et al., 1998).

[e] We assessed physical activity using the Godin Leisure-Time Exercise Questionnaire (Godin & Shephard, 1985). Both the young and older adult group medians fell within the "active" range (i.e., scores of ≥ 24). One older adult did not complete this questionnaire, so $n = 22$ older adults for this variable.

[f] Participants rated their sleepiness level from 1-7 at the start of each session using the Stanford Sleepiness Scale (Hoddes et al., 1972). One younger adult did not complete the sleepiness rating for the MRI session, so $n = 36$ young adults for this variable.

**Table A2**. Higher GSH Levels in Older Age

| | Young Adults | Older Adults | *t* | *p* | Cohen's *d* |
|---|---|---|---|---|---|
| **Frontal Voxel** | | | | | |
| CSF-Corrected GSH[a] (i.u.) | 0.90 (0.23) | 1.07 (0.29) | -2.12 | **$4.25\times10^{-2}$ *** | **0.66** |
| GM Fraction[b] | 0.49 (0.04) | 0.41 (0.04) | 8.01 | **$1.10\times10^{-9}$ ***** | **-2.33** |
| WM Fraction[b] | 0.41 (0.05) | 0.34 (0.05) | 4.85 | **$2.48\times10^{-5}$ ***** | **-1.44** |
| CSF Fraction[b] | 0.10 (0.03) | 0.25 (0.06) | -9.71 | **$2.95\times10^{-9}$ ***** | **3.51** |
| GSH Fit Error[c] (%) | 7.56 (3.24) | 8.26 (3.35) | -0.740 | 0.464 | 0.22 |
| $H_2O$ FWHM[d] (Hz) | 9.79 (1.26) | 10.56 (1.74) | -1.71 | $9.81\times10^{-2}$ | 0.55 |
| $B_0$ Frequency Offset[e] (ppm) | $3.68\times10^{-3}$ (0.01) | $1.50\times10^{-2}$ (0.01) | -3.26 | **$2.49\times10^{-3}$ **** | **0.97** |
| **Sensorimotor Voxel** | | | | | |
| CSF-Corrected GSH[a] (i.u.) | 1.04 (0.29) | 1.30 (0.46) | -2.38 | **$2.29\times10^{-2}$ *** | **0.71** |
| GM Fraction[b] | 0.51 (0.03) | 0.40 (0.04) | 11.28 | **$2.11\times10^{-14}$ ***** | **-3.17** |
| WM Fraction[b] | 0.36 (0.04) | 0.36 (0.06) | 0.06 | 0.955 | -0.02 |
| CSF Fraction[b] | 0.14 (0.05) | 0.24 (0.07) | -6.55 | **$1.14\times10^{-7}$ ***** | **1.91** |
| GSH Fit Error[c] (%) | 5.93 (1.79) | 6.71 (2.35) | -1.36 | 0.181 | 0.39 |
| $H_2O$ FWHM[d] (Hz) | 8.95 (0.80) | 9.37 (1.53) | -1.22 | 0.231 | 0.38 |
| $B_0$ Frequency Offset[e] (ppm) | $1.08\times10^{-2}$ ($9.51\times10^{-3}$) | $1.71\times10^{-2}$ ($1.13\times10^{-2}$) | -2.21 | **$3.30\times10^{-2}$ *** | **0.62** |

*Note*. *$p$ < 0.05, ***$p$ < 0.001. In the second and third columns, we report the mean ± standard deviation for each age group. In the fourth and fifth columns, we report the results of a two-sample independent t-test between groups. Each variable satisfied the normality and homogeneity of variances assumptions for a parametric two-sample t-test, with several exceptions (frontal CSF fraction, GSH fit error, and $H_2O$ FWHM; sensorimotor CSF-corrected GSH, CSF fraction, GSH fit error, and $H_2O$ FWHM). In these cases, the results of a two-sided Wilcoxon rank-sum test yielded the same interpretation as the respective parametric t-tests. In the sixth column, we report the nonparametric effect size as described by (Field et al., 2012; Rosenthal et al., 1994); see Appendix F. All significant *p*-values (bolded) remained at $p$ < 0.05 after applying the Benjamini-Hochberg FDR correction (Benjamini & Hochberg, 1995), with the exception of CSF-corrected GSH within the frontal voxel and sensorimotor $B_0$ frequency offset. These *p*-values remained only at trend-level significance after applying the correction ($p$ = 0.066 and $p$ = 0.058, respectively).

[a] Here we report the Gannet variable `GSH.ConcIU_CSFcorr`, which is the GSH concentration within the voxel, corrected for tissue composition. The Gannet variable `GSH.ConcIU` (i.e., GSH concentration, *not* accounting for age-related cortical atrophy) was not significantly different between age groups for either voxel; $p$ = 0.750 and $p$ = 0.327 for the frontal and sensorimotor voxels, respectively.

[b] Here we report the Gannet variables `fGM`, `fWM`, and `fCSF`, which indicate the fraction of gray matter (GM), white matter (WM), and cerebrospinal fluid (CSF) within the voxel.

[c] Here we report the Gannet variable `GSH.FitError_W`, which is the root sum square of the standard deviation of model residuals normalized to the model amplitude of the GSH and water signals, expressed as a percentage (Edden et al., 2014). We excluded subjects with fit error >20% prior to running any statistical tests; see Table E1.

[d] Here we report the Gannet variable `H2O.FWHM`, which is the full-width half-maximum of the unsuppressed water signal (Edden et al., 2014).

[e] Here we report the Gannet variable `AvgDeltaF0`, which is the mean difference between the observed frequency of the residual water signal in the pre-frequency-corrected subspectra and the nominal water frequency at 4.68 ppm (Mikkelsen et al., 2017).

**Table A3**. Higher GSH Levels in Sensorimotor versus Frontal Cortex

| | **Frontal-Sensorimotor Voxel Correlation** | | | | |
|---|---|---|---|---|---|
| | | ***r*** | ***p*** | ***Z*** | ***p*** |
| | Young Adults | 0.45 | **$8.96 \times 10^{-3}$ **** | 0.295 | 0.384 |
| | Older Adults | 0.52 | **$2.31 \times 10^{-2}$ *** | -- | -- |
| | **Frontal-Sensorimotor Voxel Within-Subjects Tests** | | | | |
| | | **Frontal Voxel** | **Sensorimotor Voxel** | ***t*** | ***p*** |
| CSF-Corrected GSH[a] (i.u.) | Young Adults | 0.90 (0.24) | 1.03 (0.30) | 2.55 | **0.016*** |
| | Older Adults | 1.07 (0.29) | 1.32 (0.50) | 2.57 | **0.019*** |
| GM Fraction[b] | Young Adults | 0.49 (0.04) | 0.51 (0.03) | 2.82 | **$8.14 \times 10^{-3}$**** |
| | Older Adults | 0.41 (0.04) | 0.40 (0.04) | -0.73 | 0.474 |
| WM Fraction[b] | Young Adults | 0.41 (0.05) | 0.35 (0.04) | -6.73 | **$1.33 \times 10^{-7}$***** |
| | Older Adults | 0.34 (0.05) | 0.36 (0.06) | 1.42 | 0.171 |
| CSF Fraction[b] | Young Adults | 0.10 (0.03) | 0.14 (0.05) | 6.33 | **$4.14 \times 10^{-7}$***** |
| | Older Adults | 0.25 (0.06) | 0.24 (0.07) | -0.74 | 0.467 |

*Note*. **p* < 0.05, ***p* < 0.01, ****p* < 0.001. In the top section, we report the Pearson correlation coefficient and corresponding *p*-value, as well as the results of the Fisher r-to-Z test for differences in correlation strength between groups. In the bottom section, for each age group and variable, we report the mean ± standard deviation, as well as the results of a parametric within-subjects paired t-test. All data met the appropriate assumptions for parametric t-tests, with the exception of GSH fit error and young adult CSF fraction. In these cases, the results of a paired Wilcoxon rank-sum test yielded the same interpretation as the respective parametric t-tests. GSH concentrations reported here are CSF-corrected (i.e., `GSH.ConcIU_CSFcorr`). All significant *p*-values (bolded) remained significant at *p* < 0.05 after applying the Benjamini-Hochberg FDR correction (Benjamini & Hochberg, 1995). In this table, we include only subjects who had data for both the frontal and sensorimotor voxels; see Table E1 for details.

[a] For this variable, we also conducted a between-group nonparametric two-sided Wilcoxon rank-sum test for sensorimotor GSH - frontal GSH. This was to determine whether the magnitude of the regional difference in GSH differed between age groups. This test was not significant (W = 264; *p* = 0.352).

[b] Here we report the Gannet variables `fGM`, `fWM`, and `fCSF`, which indicate the fraction of gray matter (GM), white matter (WM), and cerebrospinal fluid (CSF) within the voxel.

**Table A4**. Relationships of Frontal Voxel GSH with Cognitive and Motor Performance

| | YA Corrected Model | | | | OA Corrected Model | | | | Fisher r-to-Z Results | |
|---|---|---|---|---|---|---|---|---|---|---|
| | **GSH β (SE)** | **t** | ***p*** | **Part. r** | **GSH β (SE)** | **t** | ***p*** | **Part. r** | **Z** | ***p*** |
| **Cognition** | | | | | | | | | | |
| MoCA Score | -0.18 (1.37) | -0.13 | 0.897 | -0.02 | -0.36 (1.47) | -0.24 | 0.813 | 0.04 | 0.19 | 0.424 |
| **Balance** | | | | | | | | | | |
| A/P Sway Path | 0.17 (0.72) | 0.23 | 0.818 | 0.04 | 0.46 (0.93) | 0.50 | 0.628 | 0.13 | 0.28 | 0.391 |
| A/P Sway Speed and Variability | 0.09 (0.76) | 0.12 | 0.906 | 0.02 | -1.09 (1.05) | -1.03 | 0.317 | -0.26 | 0.93 | 0.177 |
| M/L Sway Path | 0.61 (0.74) | 0.82 | 0.421 | 0.15 | 0.18 (0.85) | 0.21 | 0.838 | 0.05 | 0.31 | 0.379 |
| M/L Sway Speed and Variability | 0.53 (0.82) | 0.64 | 0.525 | 0.12 | 0.59 (0.95) | 0.62 | 0.542 | 0.16 | 0.14 | 0.445 |
| **Gait** | | | | | | | | | | |
| **Rhythm:** Cadence (steps/min) | -2.40 (4.39) | -0.55 | 0.589 | -0.10 | 0.52 (9.05) | 0.06 | 0.955 | 0.01 | 0.37 | 0.355 |
| **Phase:** Stance (%GC) | 1.04 (1.13) | 0.92 | 0.364 | 0.17 | 0.92 (1.31) | 0.70 | 0.497 | 0.18 | 0.04 | 0.485 |
| **Pace:** Composite Score | -1.90 (1.46) | -1.30 | 0.204 | -0.23 | -1.36 (1.45) | -0.94 | 0.363 | -0.24 | 0.02 | 0.494 |
| **Variability:** Composite Score | -3.02 (1.78) | -1.69 | 0.101 | -0.30 | 2.90 (2.16) | 1.35 | 0.198 | 0.33 | **2.10** | **0.018*** |
| **Manual Dexterity** | | | | | | | | | | |
| Pegboard Composite | $-7.25\text{x}10^{-2}$ (1.22) | -0.06 | 0.953 | -0.04 | -1.46 (1.44) | -1.01 | 0.328 | -0.19 | 0.48 | 0.315 |

*Note.* $^*p < 0.05$. SE = standard error. For each age group, we present the results for the linear model testing the relationship between CSF-corrected GSH and behavioral performance, controlling for the relevant covariates (see Methods for the covariates used in each model). For ease of interpretation, we present the β, t, and *p* values for GSH only and not for the covariates. We also present the Part r. = partial r, which is the r value for the partial correlation between GSH and the performance metric, controlling for the covariates included in that model. We then present the Fisher r-to-Z results for assessing whether the strength of the partial correlation was different for the young versus older adults.

**Table A5**. Relationships of Sensorimotor Voxel GSH with Cognitive and Motor Performance

| | **YA Corrected Model** | | | | **OA Corrected Model** | | | | **Fisher *r*-to-*Z* Results** | |
|---|---|---|---|---|---|---|---|---|---|---|
| | **GSH β (SE)** | ***t*** | ***p*** | **Part. *r*** | **GSH β (SE)** | ***t*** | ***p*** | **Part. *r*** | ***Z*** | ***p*** |
| **Cognition** | | | | | | | | | | |
| MoCA Score | 0.32 (1.07) | 0.30 | 0.770 | 0.01 | -1.14 (0.96) | -1.19 | 0.250 | -0.13 | 0.50 | 0.309 |
| **Balance** | | | | | | | | | | |
| A/P Sway Path | -0.16 (0.54) | -0.30 | 0.767 | -0.05 | 0.26 (0.57) | 0.45 | 0.656 | 0.10 | 0.55 | 0.290 |
| A/P Sway Speed and Variability | -0.06 (0.58) | -0.11 | 0.913 | -0.02 | -0.28 (0.60) | -0.47 | 0.644 | -0.11 | 0.31 | 0.379 |
| M/L Sway Path | -0.22 (0.61) | -0.35 | 0.727 | -0.06 | 0.60 (0.49) | 1.23 | 0.232 | 0.27 | 1.20 | 0.115 |
| M/L Sway Speed and Variability | 0.99 (0.62) | 1.59 | 0.122 | 0.27 | 1.47 (0.44) | 3.39 | **$3.09 \times 10^{-3}$ **** | **0.61** | 1.52 | 0.064 |
| **Gait** | | | | | | | | | | |
| **Rhythm:** Cadence (steps/min) | $-1.30 \times 10^{-3}$ (3.16) | 0.00 | 1.000 | < 0.01 | -7.53 (4.85) | -1.55 | 0.137 | -0.34 | 1.22 | 0.111 |
| **Phase:** Stance (%GC) | -0.22 (0.88) | -0.25 | 0.806 | -0.04 | 0.62 (0.83) | 0.74 | 0.468 | 0.17 | 0.75 | 0.227 |
| **Pace:** Composite Score | -0.12 (1.11) | -0.11 | 0.917 | -0.02 | -0.55 (0.95) | -0.58 | 0.566 | -0.13 | 0.40 | 0.344 |
| **Variability:** Composite Score | -1.30 (1.40) | -0.93 | 0.360 | -0.16 | 3.85 (1.00) | 3.83 | **$1.12 \times 10^{-3}$ **** | **0.66** | 3.37 | **$4.00 \times 10^{-4}$ ***** |
| **Manual Dexterity** | | | | | | | | | | |
| Pegboard Composite | 0.25 (0.99) | 0.26 | 0.798 | 0.03 | -2.51 (0.63) | -3.97 | **$7.58 \times 10^{-4}$ ***** | **-0.57** | 2.40 | **0.008**** |

*Note.* ***p* < 0.01, ****p* < 0.001. SE = standard error. For each age group, we present the results for the linear model testing the relationship between CSF-corrected GSH and behavioral performance, controlling for the relevant covariates (see Methods for the covariates used in each model). For ease of interpretation, we present the β, t, and *p* values for GSH only and not for the covariates. We also present the Part r. = partial r, which is the r value for the partial correlation between GSH and the performance metric, controlling for the covariates included in that model. We then present the Fisher r-to-Z test results for assessing whether the strength of the partial correlation was different for the young versus older adults. All significant *p*-values (bolded) remained significant at $p<0.05$ after applying the Benjamini-Hochberg FDR correction (Benjamini & Hochberg, 1995). Moreover, although for ease of interpretation we present here the results of linear models calculated using `lm`, in all cases where the *p* value was significant in the parametric linear model, the corresponding *p* value was also significant for the analogous nonparametric model.

### **Appendix B:** Selection and Analysis of Balance Variables

Participants completed a static balance task (i.e., standing as still as possible for 30 seconds) under four conditions: eyes open (EO), eyes closed (EC), eyes open-foam (EOF), and eyes closed-foam (ECF). We selected only the eyes open condition for further analysis because there are age differences in postural sway during quiet stance with the eyes open (Baloh et al., 1994; Maki et al., 1990), and eyes open postural sway is predictive of falls for older adults (Fernie et al., 1982; Maki et al., 1990). Additionally, both foam conditions had several clear outliers (EOF: 1 young outlier ~5 standard deviations (SDs) greater than the group mean for multiple measures, ECF: 1 young and 2 older outliers ~3 SDs greater than the group mean for multiple measures). Conversely, using the EO condition allowed us to include all participants in subsequent analyses.

We extracted the 25 “acceleration” postural sway variables from MobilityLab for the EO condition. To reduce the number of variables, we conducted an exploratory factor analysis (EFA). The variables met the EFA assumptions of linearity, interval data, no outliers, and a sufficiently large sample size. Several variables did not meet the EFA assumption of perfect multicollinearity. We checked this assumption by calculating a correlation matrix between all 25 variables (Fig. B1). Based on this correlation matrix, before running the EFA, we removed five variables that had perfect multicollinearity with another variable ($r = 0.98$-$1.0$): jerk, mean velocity, RMS sway, 95% ellipse axis 2 radius, and range.

We then used `fa.parallel` in the `psych` package (Revelle, 2014) to produce a scree plot, which suggested four factors as the optimal number for these data. Next, we conducted an EFA using the `fa` function (Revelle, 2014); we asked for four factors, oblique rotation, maximum likelihood estimation, and the Bartlett method for computation of factor scores. Factor loadings are presented in Table B1, and visualization of the factors is presented in Fig. B2. Based on the loadings ≥ 0.5, we named the four factors as follows: anterior/posterior (A/P) sway path, A/P sway speed and variability, medial/lateral (M/L) sway path, and M/L sway speed and variability. Together, these four factors explained a cumulative variance of 75% in the data.

We then extracted composite scores with the Bartlett method (Bartlett, 1937) for use in subsequent analyses. The Bartlett method produces unbiased standard score estimates (mean = 0, SD = the squared multiple correlation between the items and factor) (DiStefano et al., 2009). We selected to use the Bartlett method for several reasons: 1) the Bartlett method produces high validity, unbiased estimates between each factor score and factor (DiStefano et al., 2009); 2) Bartlett method scores are more likely to represent the true factor scores compared to other factor score estimation methods (e.g., regression or Anderson-Rubin) (DiStefano et al., 2009); and 3) estimated factor scores (i.e., using a method such as Bartlett) provide greater precision than simple sums based on an arbitrary cut-off value or weighted sum composite scores based on the factor loadings (DiStefano et al., 2009).

**Table B1.** Factor Loadings for Balance Variables

| Variable | Variable Description[a] | ML1: A/P Sway Path | ML3: A/P Sway Speed and Variability | ML4: M/L Sway Speed and Variability | ML2: M/L Sway Path |
|---|---|---|---|---|---|
| Path Length, Sagittal Plane ($m/s^2$) | Total length of the sway path in the M/L direction. | 0.936 | | | |
| Jerk, Sagittal Plane ($m^2/s^5$) | Smoothness of sway from the time derivative of the sway path in the A/P direction. | 0.908 | | | |
| Path Length ($m/s^2$) | Sway path, total length of COP (acceleration) trajectory | 0.737 | | | |
| Frequency Dispersion (AD) | Frequency dispersion in the transverse plane. | -0.552 | | | |
| Mean Velocity, Sagittal Plane (m/s) | Mean velocity of the sway path in the A/P direction. | | 0.907 | | |
| RMS Sway, Sagittal Plane ($m/s^2$) | The root mean square of the sway angle in the sagittal plane. | | 0.804 | | |
| Range, Sagittal Plane ($m/s^2$) | Total range of the sway path in the A/P direction. | | 0.771 | | |
| Centroidal Frequency (Hz) | Frequency of sway from the centroid of the sway path's power spectrum in the transverse plane. | | -0.760 | | |
| 95% Ellipse Axis 1 Radius ($m^2/s^2$) | The minor axis of the ellipse that is best fit to the sway, using a 95% confidence interval. | | | 0.895 | |
| RMS Sway, Coronal Plane ($m/s^2$) | The root mean square of the sway angle in the coronal plane. | | | 0.706 | |
| Range, Coronal Plane ($m/s^2$) | Total range of the sway path in the M/L direction. | | | 0.664 | |
| Mean Velocity, Coronal Plane (m/s) | Mean velocity of the sway path in the M/L direction. | | | 0.569 | |
| 95% Ellipse Sway Area | The area of an ellipse covering 95% of the sway angle in the | | | 0.569 | |

| | | | | | |
|---|---|---|---|---|---|
| $(m^2/s^4)$ | coronal and sagittal planes. | | | | |
| Path Length, Coronal Plane $(m/s^2)$ | Sway path, total length of COP (acceleration) trajectory | | | | 0.891 |
| Jerk, Coronal Plane $(m^2/s^5)$ | Smoothness of sway from the time derivative of the sway path in the M/L direction. | | | | 0.808 |
| Centroidal Frequency, Coronal Plane (Hz) | Frequency of sway from the centroid of the sway path's power spectrum in the M/L direction. | | | | 0.721 |
| Frequency Dispersion, Coronal Plane (AD) | Frequency dispersion in the M/L direction. | | | 0.544 | -0.619 |
| Frequency Dispersion, Sagittal Plane (AD) | Frequency dispersion in the A/P direction. | | | | |
| Centroidal Frequency, Sagittal Plane (Hz) | Frequency of sway from the centroid of the sway path's power spectrum in the A/P direction. | | | | |
| 95% Ellipse Rotation (radians) | The counterclockwise rotation of the best fit ellipse, where no rotation results in the major axis of the ellipse being aligned with the coronal plane. | | | | |

*Note.* M/L = medial/lateral; A/P = anterior/posterior. To increase interpretability, in this table we include only the factor loadings ≥ 0.5. Three variables (listed at the bottom of the table) did not load onto any factors at ≥ 0.5.

[a] These variable descriptions were obtained from the APDM MobilityLab support website: https://support.apdm.com/en-us/article_attachments/Metric_Definitions/

**Fig. B1.** Correlation Matrix of All Balance Variables

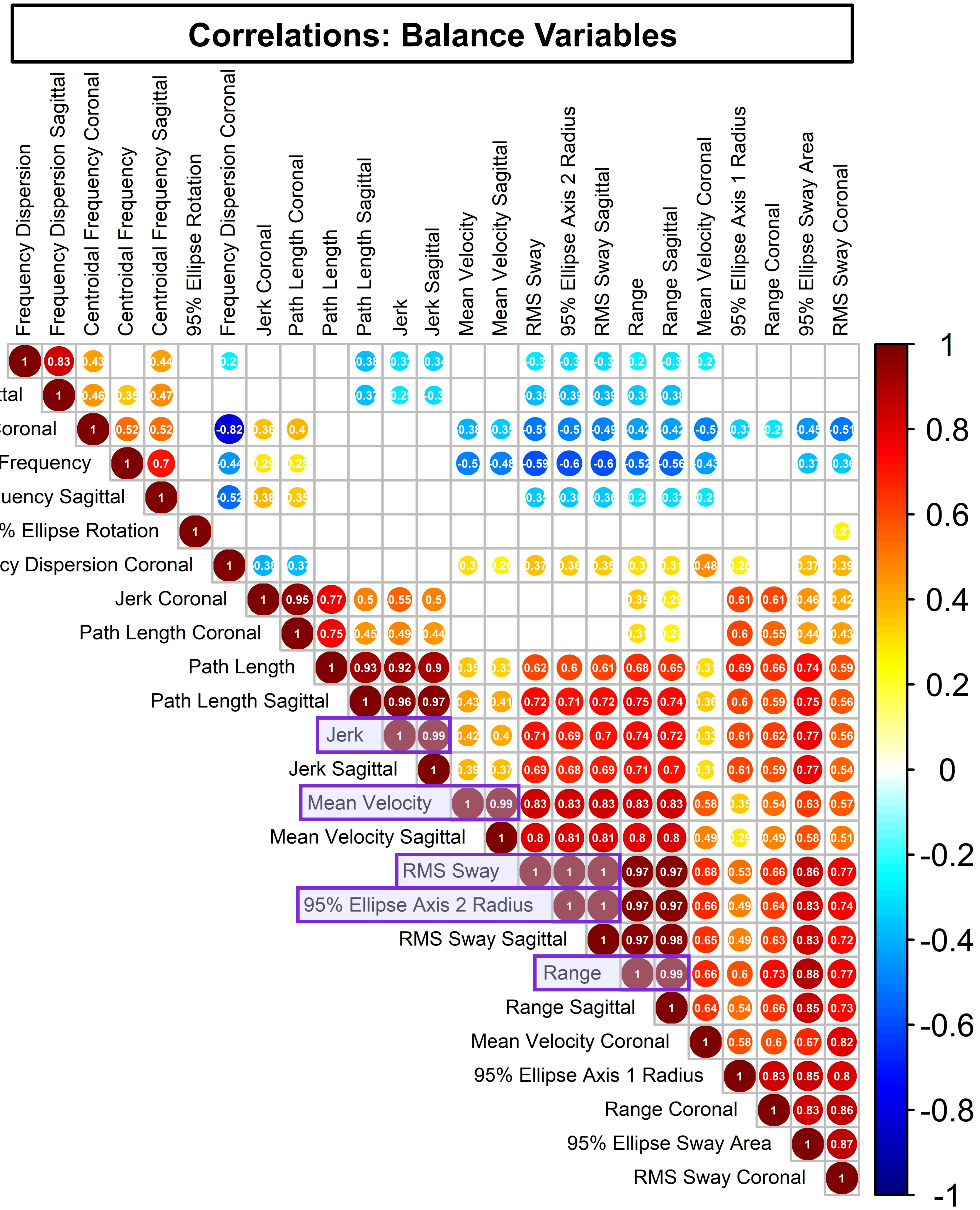


*Note.* Here we present the Pearson correlation matrix for all of the gait variables, created using `corrplot` in the `corrplot` package (Wei et al., 2017). Circle size represents the strength of the correlation, and correlation coefficient values are printed in white. Warm colors represent positive correlation coefficient values, and cool colors represent negative correlation coefficient values. This plot includes each of the 25 sway variables used to test the perfect multicollinearity assumption. The five variables indicated with a purple box were removed before conducting the EFA due to perfect multicollinearity (i.e., r = 0.98-1.0).

**Fig. B2.** Balance Factors

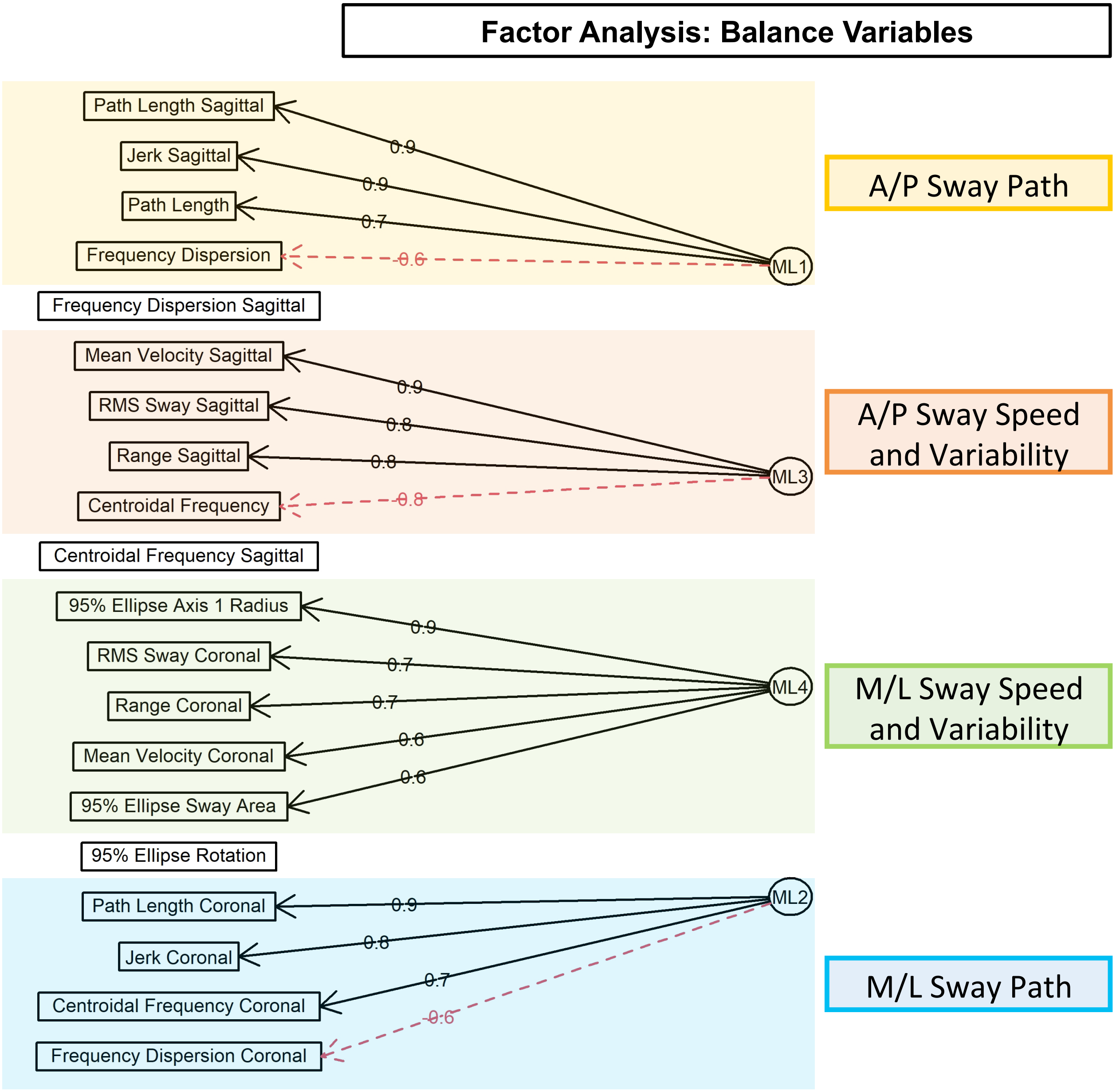


*Note*. Here we present the results of the balance variable EFA. We created this plot using `fa.diagram` in the `psych` package (Revelle, 2014). This plot includes the factor loadings for the 20 sway variables included in the EFA. For ease of interpretation, only factor loadings ≥ 0.5 are shown here. For the one variable that loaded onto two factors (i.e., frequency dispersion, coronal), only the higher factor loading is pictured here (i.e., the loading of this variable onto ML2). We do not depict a relationship with any of the four factors for the three variables that did not load onto any factors at ≥ 0.5

### **Appendix C:** Selection and Analysis of Gait Variables

For the 4-minute walk, we extracted 18 left- and right-sided spatiotemporal gait variables from MobilityLab. Next, we calculated the mean of the left- and right-sided variables to obtain 9 average variables (Table C1). We also used MobilityLab output to calculate 10 metrics of gait variability—for stride length, stride time, step time, swing %, and stance %. We calculated the standard deviation (SD) across all gait cycles for the left and right side for these metrics. We then calculated the mean of the left- and right-sided SDs to obtain 5 average variability metrics (Table C1).

Next, we computed the correlation matrix (Fig. C1) between these 14 average variables to determine whether we could implement an exploratory factor analysis (EFA) to obtain gait summary metrics, as we did for the balance variables. However, multiple variables exhibited perfect multicollinearity (e.g., swing % is the exact inverse of stance %). In all cases of perfect multicollinearity (i.e., correlation coefficient r = 0.98-1.0), we selected only one variable from each pair for further analysis. After these exclusions, too few variables remained to conduct an EFA.

Thus, instead of an EFA, we categorized the remaining gait variables based on the divisions described in (Hollman et al., 2011). Hollman and colleagues measured overground gait at a self-selected pace for 294 older adults aged 70+ years. They extracted 23 spatiotemporal gait parameters and conducted an EFA. Their EFA yielded five "domains" of gait. We used this structure to sort our gait variables into four domains: rhythm, phase, pace, and variability (Figs. C1-C2). Of note, MobilityLab cannot calculate step width, so we were unable to use Hollman and colleagues' fifth domain, base of support.

We defined scores for our four gait domains as follows:

1. **Rhythm:** We selected cadence (steps/min) to represent the rhythm domain and removed the two other rhythm variables (i.e., step time and stride time) from further analyses, as these metrics correlated perfectly (r = -0.99) with cadence.

2. **Phase:** We selected stance (% gait cycle) to represent the phase domain and removed the other three phase variables (i.e., swing %, double support %, and single support %) from further analyses, as these metrics correlated perfectly (r = -1, 1, and -1, respectively) with stance %.

3. **Pace:** For the pace domain, the two variables (i.e., walking speed and stride length) did *not* correlate perfectly (r = 0.84). Thus, we computed a composite score for pace. We converted the two pace variables to standardized Z scores using the formula (data point - mean)/(SD). We used the mean and SD for the entire sample (rather than for the two age groups separately). Next, we summed these Z scores to calculate a composite pace score. These two variables were *positively* correlated, so a simple sum was appropriate for the creation of a composite score.

4. **Variability:** For the variability domain, we removed the two metrics that correlated perfectly with another variable (i.e., stance % variability and stride time variability, r = 0.98-1). This left us with three variables that did *not* correlate perfectly (r = 0.34-0.61). We converted these three variables to standardized Z scores, as described above, and summed these Z scores to obtain a composite variability score. Each of these variables was *positively* correlated, so a simple sum was appropriate for the creation of a composite score.

**Table C1.** Gait Variables

| Variable | Unit | Variable Description[a] |
|---|---|---|
| **Gait Rhythm** | | |
| Cadence[b] | steps/min | The number of steps per minute, counting steps made by both feet |
| Step Time | s | The duration of a step, measured as the period from initial contact of one foot to the next initial contact of the opposite foot |
| Stride Time | s | The duration of a full gait cycle, measured from the left foot's initial contact to the next initial contact of the left foot |
| **Gait Phase** | | |
| Stance %[b] | % Gait Cycle | The percentage of the gait cycle in which the foot is on the ground |
| Swing % | % Gait Cycle | The percentage of the gait cycle in which the foot is not on the ground |
| Single Support % | % Gait Cycle | The percentage of the gait cycle in which the opposite foot is not touching the ground |
| Double Support % | % Gait Cycle | The percentage of the gait cycle in which both feet are on the ground |
| **Gait Pace** | | |
| Walk Speed | m/s | The forward speed of the subject, measured as the forward distance traveled during the gait cycle divided by the gait cycle duration |
| Stride Length | m | The forward distance travelled by the foot during a gait cycle |
| **Gait Variability** | | |
| Step Time Variability[c] | SD | Average of left and right step time SD |
| Stride Time Variability | SD | Average of left and right stride time SD |
| Swing % Variability[c] | SD | Average of left and right swing % SD |
| Stance % Variability | SD | Average of left and right stance % SD |
| Stride Length Variability[c] | SD | Average of left and right stride length SD |

*Note*. s = seconds; m = meters; SD = standard deviation.

[a] Variable descriptions were obtained from the MobilityLab support website: https://support.apdm.com/en-us/article_attachments/Metric_Definitions/

[b] For these domains, due to instances of perfect multicollinearity, we used *only* the variable indicated with the superscript ‘b’ to represent the respective gait domain.

[c] For gait variability, we used *only* the variables indicated with the superscript ‘c’ to create a composite score. The other two variables were removed due to perfect multicollinearity with one or more other variables.

**Fig. C1.** Correlation Matrix of All Gait Variables

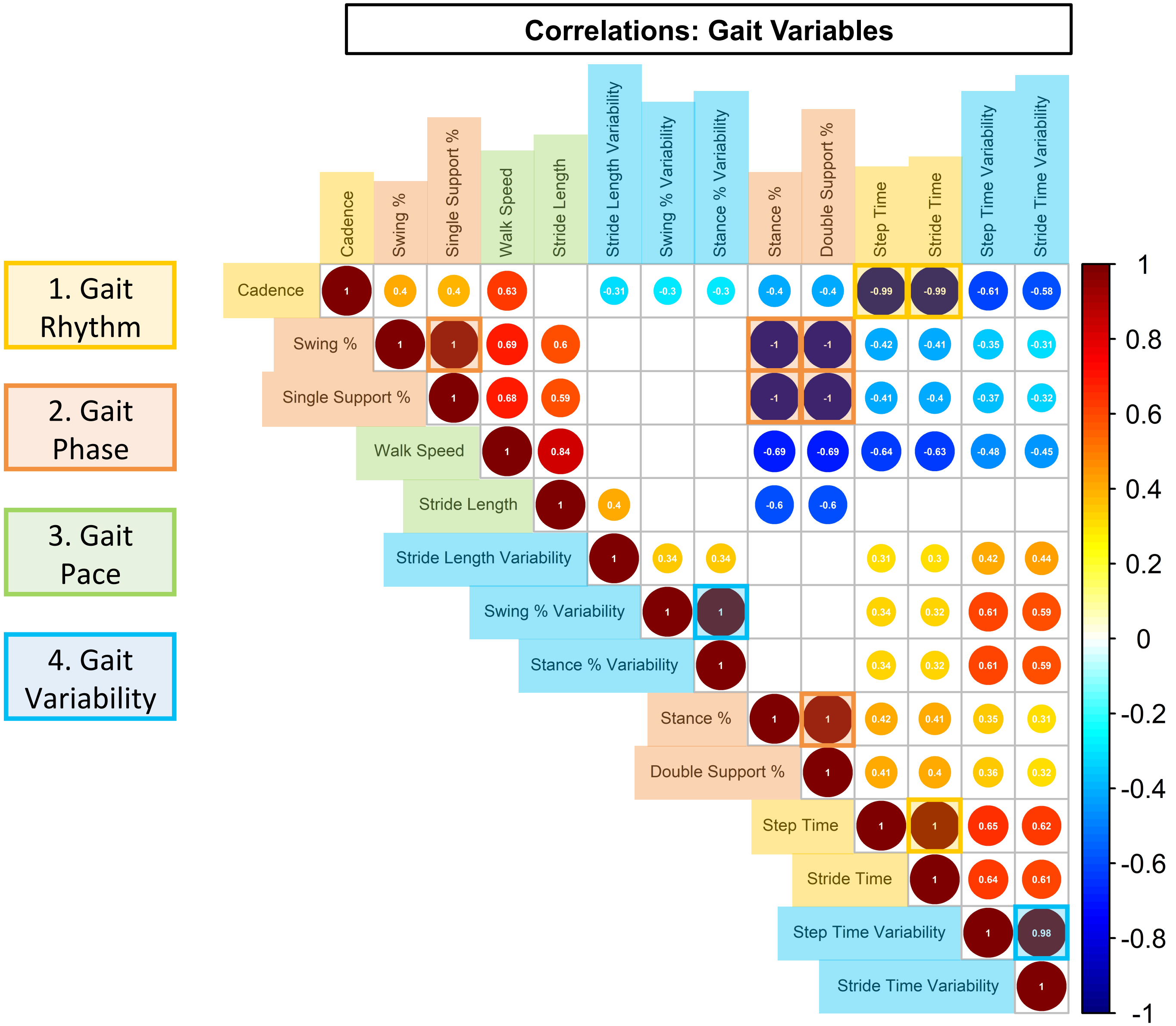


*Note.* Here we present the Pearson correlation matrix for all of the gait variables, created using `corrplot` in the `corrplot` package (Wei et al., 2017). Circle size represents the strength of the correlation, and correlation coefficient values are printed in white. Warm colors represent positive correlation coefficient values, and cool colors represent negative correlation coefficient values. The four gait domains are presented to the left of the correlation matrix. Each gait domain and its corresponding variables are highlighted in the same color. The colored boxes within the correlation matrix highlight any instances of perfect multicollinearity. In such cases of perfect multicollinearity, we selected only one variable from the correlated pair to include in further analyses; see text above for details.

**Fig. C2.** Summary of Gait Variables Used in Further Analyses

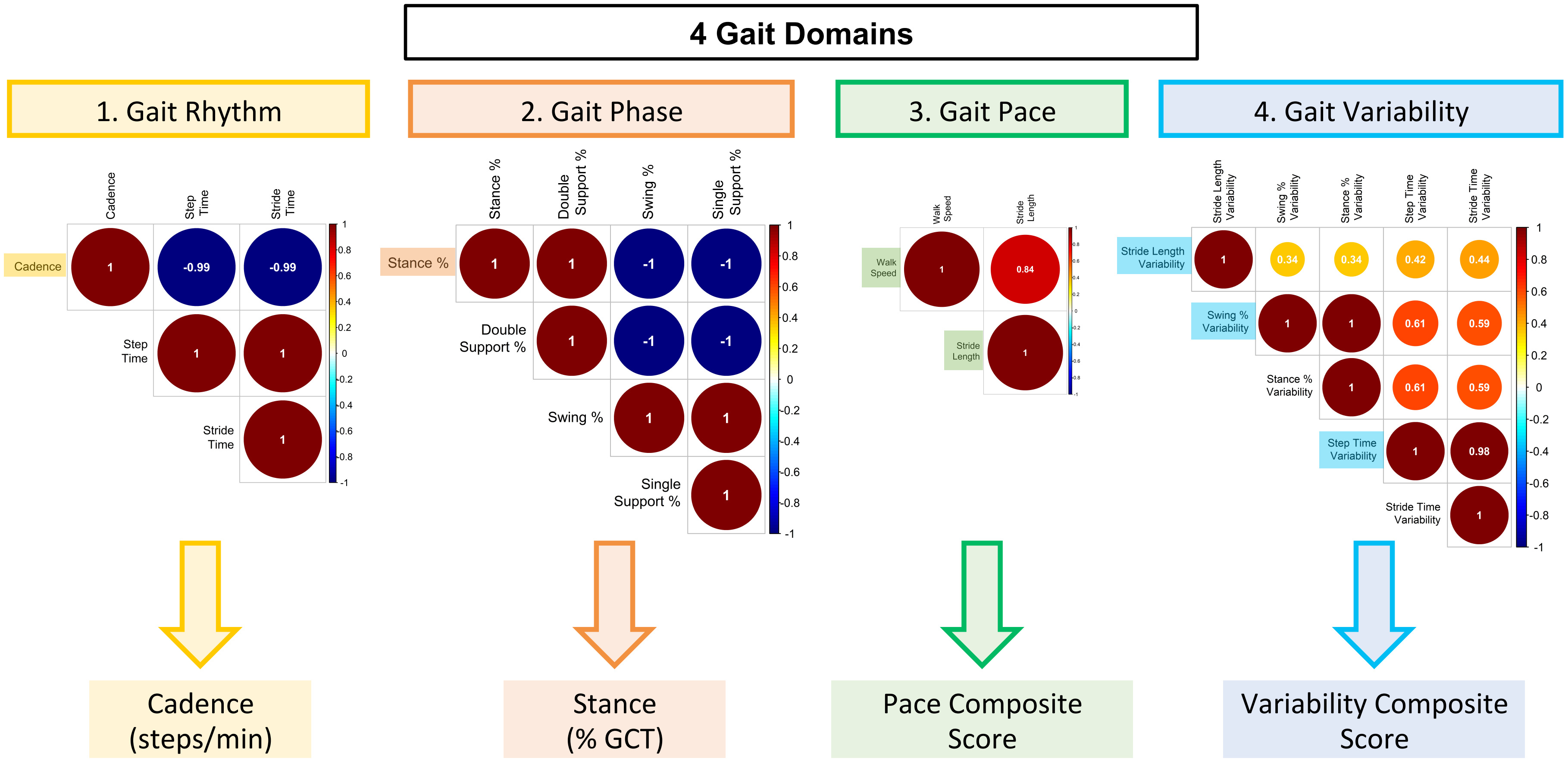


*Note.* Here we present the Pearson correlation matrix, created using `corrplot` in the `corrplot` package (Wei et al., 2017), for all of the gait variables, separated into the four gait domains. Circle size represents the strength of the correlation, and correlation coefficient values are printed in white. Warm colors represent positive correlation coefficient values, and cool colors represent negative correlation coefficient values. In each correlation matrix, we highlight the names of the variables used in further analyses (i.e., either as the representative variable for that domain or as part of a composite score for that domain). We indicate below each matrix whether one variable was selected to represent that gait domain (due to perfect multicollinearity, in the cases of gait rhythm and phase), or if a composite score was created

### **Appendix D:** Age Differences in Cognitive and Motor Performance

For completeness, here we report tests of age differences in performance of the cognitive and motor tasks. There were several age differences in these behavioral measures (Table D1). Older adults had lower MoCA scores. No age differences emerged for the balance metrics, except for M/L sway path, on which older adults scored better (i.e., *less* postural sway) than young adults, although the effect size for this group difference was small (-0.30). No age differences emerged for the gait metrics, except for the phase domain. Older adults spent *longer* in the stance phase than younger adults. Additionally, older adults had *poorer* pegboard scores compared to young adults.

**Table D1.** Age Differences in Cognitive and Motor Performance

| | Young Adults | Older Adults | W[a] | *p* | Effect Size |
|---|---|---|---|---|---|
| **Cognition** | | | | | |
| MoCA Score | 28 (3) | 27 (2) | 580.5 | **0.017*** | **-0.31** |
| **Balance** | | | | | |
| A/P Sway Path | -0.28 (0.69) | 0.16 (1.17) | 321 | 0.114 | -0.20 |
| A/P Sway Speed and Variability | -0.51 (1.01) | -0.0043 (0.90) | 311 | 0.083 | -0.22 |
| M/L Sway Path | 0.10 (1.1) | -0.61 (1.24) | 579 | **0.020*** | **-0.30** |
| M/L Sway Speed and Variability | -0.36 (0.92) | -0.16 (1.64) | 422 | 0.964 | -0.01 |
| **Gait** | | | | | |
| **Rhythm:** Cadence (steps/min) | 104.7 (6.73) | 108.08 (16.40) | 349 | 0.248 | -0.15 |
| **Phase:** Stance (%GC) | 59.44 (1.61) | 61.26 (1.46) | 227 | **0.003**** | **-0.39** |
| **Pace:** Composite Score | -0.11 (1.85) | -0.60 (1.44) | 545 | 0.070 | -0.23 |
| **Variability:** Composite Score | -0.02 (3.90) | -0.67 (2.24) | 477 | 0.438 | -0.10 |
| **Manual Dexterity** | | | | | |
| Pegboard Composite | 1.10 (2.02) | -1.80 (1.21) | 799.5 | **$1.30 \times 10^{-8}$ ***** | **-0.73** |

*Note*. **p* < 0.05, ***p* < 0.01, ****p* < 0.001. All subjects (37 young and 23 older adults) are included in the comparisons in this table. However, we excluded several individuals from the subsequent analyses testing for GSH relationships with these behavioral metrics; see Table E1. In the second and third columns, we report the median ± interquartile range (IQR) for each age group. In the fourth and fifth columns, we report the results of nonparametric two-sample, two-sided Wilcoxon rank-sum tests. In the sixth column, we report the nonparametric effect size as described by (Field et al., 2012; Rosenthal et al., 1994); see Appendix F. All significant *p*-values (bolded) remained significant at *p* < 0.05 after applying the Benjamini-Hochberg FDR correction (Benjamini & Hochberg, 1995).

### **Appendix E:** MRS Exclusions and Quality Control

**Table E1**. Exclusions and Sample Sizes

| ***Frontal Voxel*** | | | | | | |
|---|---|---|---|---|---|---|
| | Young Adults | | | Older Adults | | |
| | Subj. ID | Reason for Exclusion | Total *n* | Subj. ID | Reason for Exclusion | Total *n* |
| Frontal Voxel GSH[a] | 1030 | GSH fit error ≥ 20% | 34 | 2003 | GSH fit error ≥ 20% | 19 |
| | 1037 | GSH fit error ≥ 20% | | 2004 | RobustSpecReg failed | |
| | 1035 | RobustSpecReg failed | | 2014 | RobustSpecReg failed | |
| | | | | 2029 | RobustSpecReg failed | |

| ***Sensorimotor Voxel*** | | | | | | |
|---|---|---|---|---|---|---|
| | Young Adults | | | Older Adults | | |
| | Subj. ID | Reason for Exclusion | Total *n* | Subj. ID | Reason for Exclusion | Total *n* |
| Sensorimotor Voxel GSH[a] | 1029 | GSH fit error ≥ 20% | 35 | -- | *No exclusions* | 23 |
| | 1035 | RobustSpecReg failed | | | | |

[a] For all analyses involving GSH, we excluded datasets if the Gannet RobustSpecReg procedure failed to provide appropriate registration of the spectra. We also excluded datasets with `GSH.FitError_W` ≥ 20%, which equated to approximately 2.5 standard deviations (SDs) above the group mean fit error for GSH.

**Fig. E1**. Age group differences in GSH, indicating subject collected with 20-channel head coil instead of 64-channel head coil

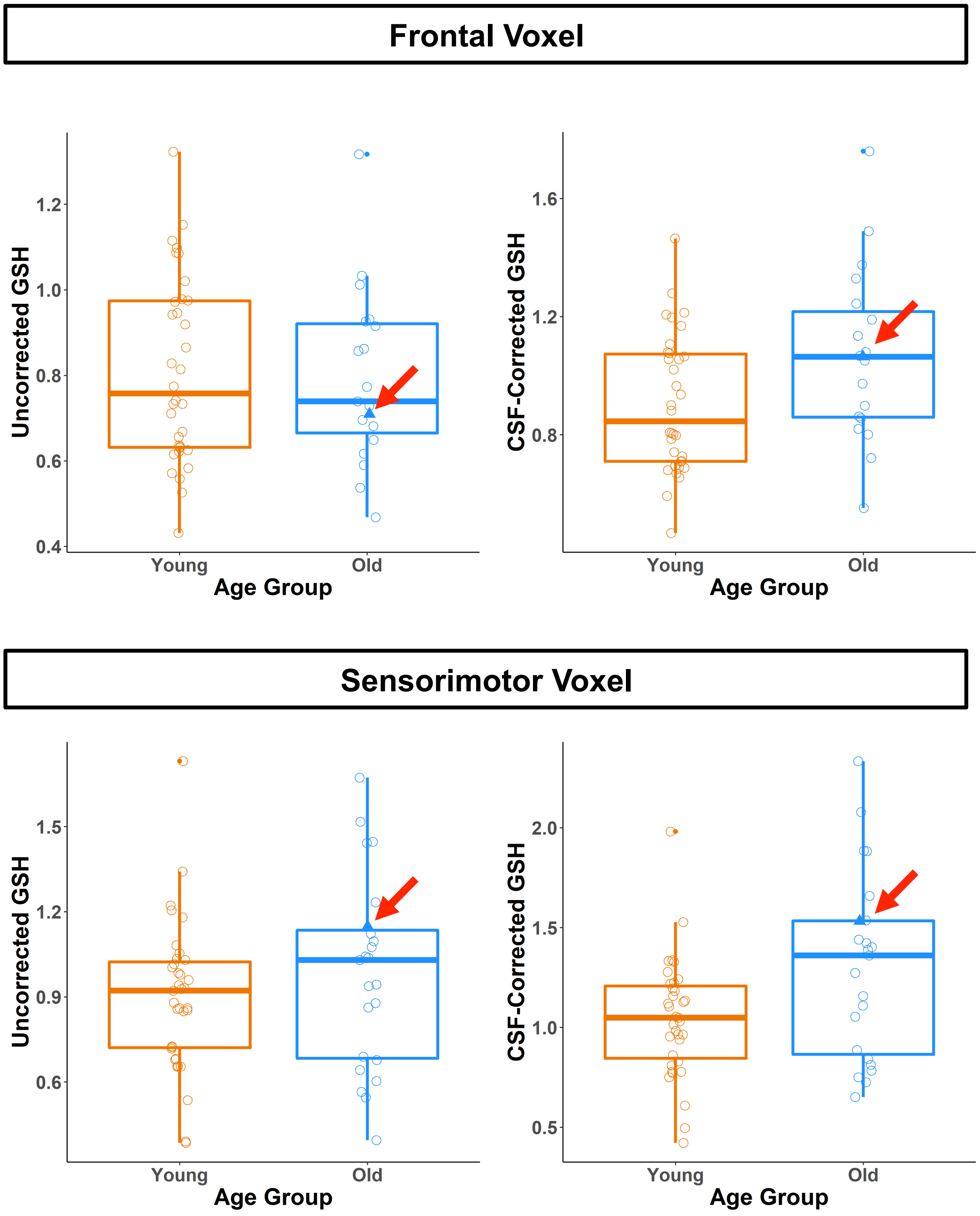


*Note.* Circles indicate young (orange) and older (blue) adults. The filled blue triangles indicate the older adult for whom data was collected with a 20-channel MRI coil, rather than the 64-channel coil used for all other subjects (due to his large head size). GSH values for this subject fell within a reasonable range of that of the other older subjects and thus were not excluded.

### **Appendix F:** Calculation of Nonparametric Effect Sizes

In order to calculate nonparametric effect sizes, we used the following formula, as described in: (Field et al., 2012; Rosenthal et al., 1994). We also thank Vincente Esparza Villalpando for providing this code; for details, see: https://www.researchgate.net/post/How_can_I_calculate_the_effect_size_for_Wilcoxon_signed_rank_test.

```
FromWilcox <- function(wilcoxModel,N)
      {
        z<-qnorm(wilcoxModel$p.value/2)
        r<-z/sqrt(N)
        cat(wilcoxModel$data.name,"Effect Size, r=",r)
      }
```
